# RhlR quorum-sensing receptor ligand sensitivity regulates the differential expression of phenazine genes in *Pseudomonas aeruginosa*

**DOI:** 10.64898/2025.12.08.692981

**Authors:** Autumn N. Pope, Varun R. Bavda, Megan L. Schumacher, Alicia G. Mendoza, Anne M. Stringer, Caleb P. Mallery, Anna Czachor, Amanda F. Kurtz, Biqing Liang, Xia Ke, Joseph T. Wade, Jon E. Paczkowski

**Author notes:** Address correspondence to Jon E. Paczkowski. Department of Medicine, New York University, Grossman School of Medicine, New York, NY, USA.

## Abstract

Bacteria control individualistic and group behaviors using a form of cell-cell communication called quorum sensing. Quorum sensing relies on the production of chemical signals called autoinducers and the subsequent detection of those signals by a cognate receptor. Many Gram-negative bacteria use the LuxR-type family of transcription factor receptors that bind to acyl-homoserine lactone autoinducer signals to regulate their function as DNA-binding proteins. A subclass of this family of transcription factor receptors requires their cognate autoinducer to fold and dimerize to bind DNA to regulate gene expression and, thus, traits associated with quorum sensing, such as biofilm formation and virulence factor production. Here, we use a chemical-genetic approach to determine the structural basis for ligand selection by the quorum-sensing receptor RhlR from *Pseudomonas aeruginosa*. The native ligand for RhlR is *N*-butyryl-L-homoserine lactone, and this protein-ligand interaction is important for initiating gene expression in *P. aeruginosa*. We determine key residues that drive ligand specificity and selectivity of RhlR to define the role of ligand-driven RhlR-dependent gene regulation of quorum-sensing traits, namely the differential expression of the phenazine genes, which encode the enzymes responsible for the synthesis of the redox-sensitive virulence factor pyocyanin, among other phenazines. Furthermore, we provide a chemical-genetic framework for future studies aimed at disrupting the RhlR-ligand interaction to suppress virulence in *P. aeruginosa*, an important nosocomial pathogen with widespread antimicrobial resistance.

## INTRODUCTION

*Pseudomonas aeruginosa* is an opportunistic pathogen that causes thousands of deaths every year in the United States (1). Virulence and pathogenesis are largely mediated by quorum sensing (QS), a cell-cell communication mechanism that relies on the production, detection, and group-wide response to signaling molecules called autoinducers (AI) (2). Typically, an AI binds to its cognate receptor to initiate changes in gene expression, often in a positive feedback manner, to alter community behavior. *P. aeruginosa*, and many other Gram-negative bacteria, use LuxR-type receptors to detect diffusible *N*-acylhomoserine lactone (AHL) AI (3) (Figure 1A). LuxR-type receptors are a class of transcription factor receptors with a highly variable N-terminal ligand binding domain (LBD) and a well-conserved C-terminal helix-turn-helix (HTH) DNA binding domain (DBD). Within the class of LuxR-type receptors there exist three different well-characterized subclasses: 1) those that require AHL to fold, dimerize, and bind DNA to act as transcriptional activators; 2) those that can fold and dimerize in the absence of the AHL, but require free exchange of AHL to bind DNA to act as transcriptional activators; 3) those that can fold, dimerize, and bind DNA to act as transcriptional repressors only in the absence of AHL and whose repression is relieved by binding to AHL (3–15). To date, three LuxR-type receptors have been characterized in *P. aeruginosa*: LasR, RhlR, and QscR (7,16–18). LasR and RhlR bind the AHLs *N*-(3-oxododecanoyl)-L-homoserine lactone (3OC_12_HSL) and *N*-butyryl-L-homoserine lactone (C_4_HSL), respectively (6,17). 3OC_12_HSL and C_4_HSL are synthesized by the LuxI synthases LasI and RhlI, respectively. Thus, LasR and RhlR have partner synthases that produce a dedicated AHL. In each case, the receptor upregulates the expression of the partner synthase gene, initiating a positive feedback loop signaling mechanism to enhance QS traits (19–21). Conversely, QscR is an orphan receptor (*i.e.*, lacks a partner synthase) and binds to the LasI-produced 3OC_12_HSL as well as other AHLs to regulate its activity (22). Each of the three receptors co-regulate a shared regulon as well as their own set of genes (8,23–26).

**Figure 1.**
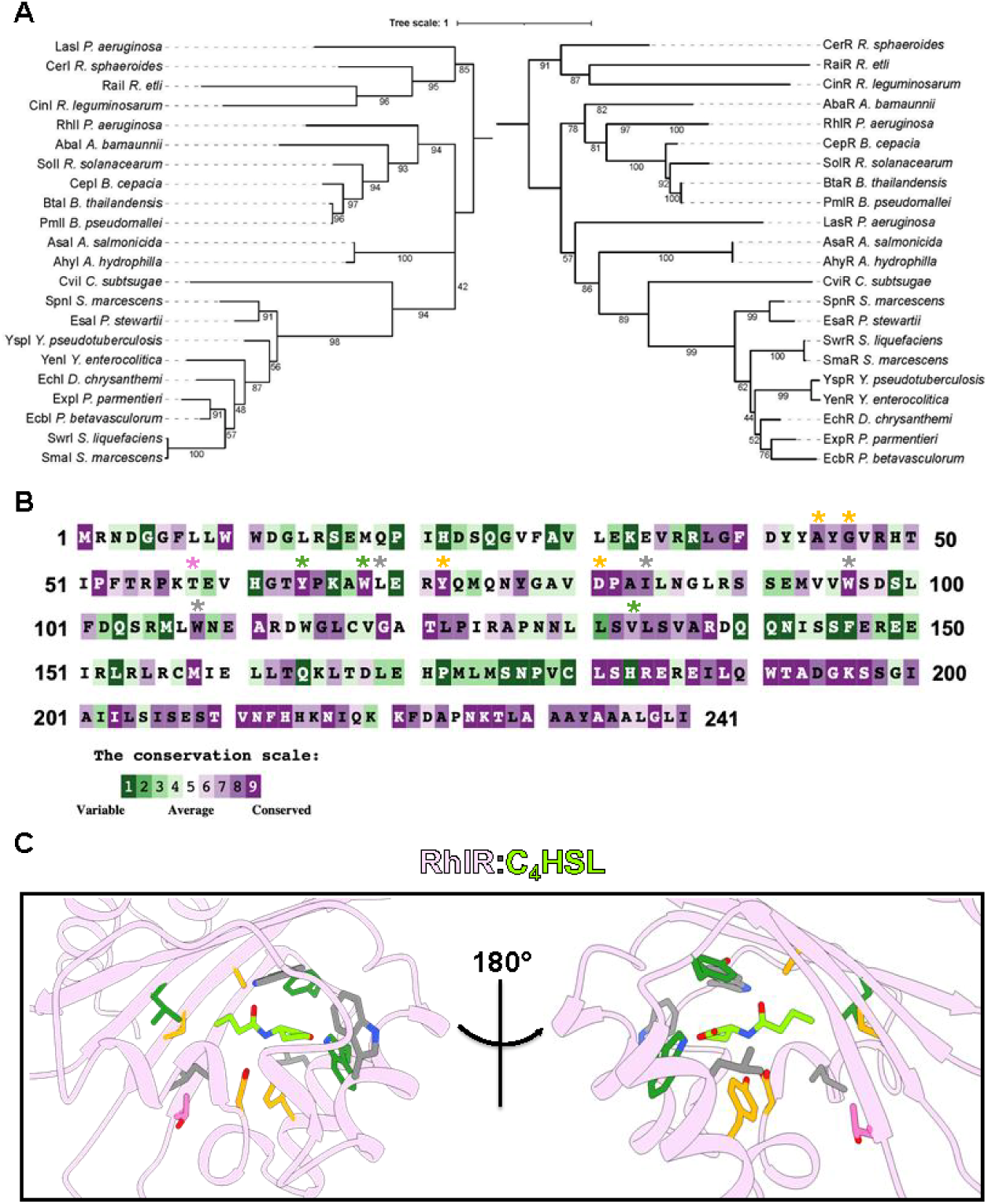
Evolutionary and structural conservation of LuxR-type transcription factor receptors. **A)** Maximum-likelihood phylogenetic trees of RhlR and RhlI homologs. The trees were constructed based on the alignment of 22 amino acid sequences for RhlI (left) and RhlR (right) homologs from well-characterized LuxIR-type systems. Bootstrap support values, indicated at the nodes, were derived from 1,000 replicates. Species names are abbreviated, and protein names are indicated along each branch. **B)** Conservation analysis of the RhlR protein sequence. The amino acid sequence of RhlR is colored based on conservation scores estimated using the ConsurfDB, with 300 homologs used in the analysis. Purple residues are highly conserved, and green residues are highly variable. Residues A44, G46, T58, Y64, W68, L69, Y72, D81, I84, W96, W108, and V133 are marked with asterisks and colored based on variant responses to C_4_HSL: orange = substitutions are recalcitrant to C_4_HSL; magenta = substitutions are hypersensitive to C_4_HSL; gray = substitutions are hyposensitive to C_4_HSL; dark green = substitutions were previously described and lead to a “constitutive” state for RhlR. **C)** Zoom in view of the RhlR (pink) LBP. C_4_HSL (green) was computationally docked into the WT RhlR LBP (PDB: 8DQ0). Residues A44, G46, T58, Y64, W68, L69, Y72, D81, I84, W96, W108, and V133 are shown as sticks. Residues are colored as in (**B**).

LuxR-type receptors rely on a bipartite ligand binding pocket (LBP) to accommodate the hydrophilic lactone head group and the hydrophobic acyl-acyl tail. Specifically, two universally conserved residues comprising a YXXXW motif are responsible for coordinating the lactone head group (Figure 1B) (27–29). The residues responsible for binding the acyl-acyl tail are divergent among LuxR-type homologs, with the size of the hydrophobic side chains dictating the acyl chain length and the pose of the tail within the LBP (Figure 1B). For example, LasR A127 is responsible for accommodating the C_12_ acyl tail group, one of the longer acyl chain groups known to be naturally synthesized by LuxI synthases. Substitution of A127 to a tryptophan resulted in a complete loss of activation by 3OC_12_HSL but improved the ability of the LasR A127W variant to respond to the non-cognate ligand 3OC_10_HSL compared to WT LasR (28). Conversely, SdiA, a LuxR-type protein from *Escherichia coli*, possesses a phenylalanine at the same site in the structure as LasR A127, and this results in 3OC_8_HSL being the ligand with the highest binding affinity (28). A chemical-genetic approach using AI analogs revealed that the LasR binding site is highly malleable, allowing for multiple conformations within the LBP with different molecule orientations driving different efficacies in WT LasR (27). Specifically, residue T75 in LasR can contact the lactone head group of certain ligands to lock them in the LBP, leading to a change in conformation of the tail group, which resulted in a molecule with low efficacy. Conversely, a LasR T75V variant reversed the binding modality of the low-affinity ligand, resulting in a rotation of the lactone head group to the canonical form and a reorientation of the tail group, which coincided with increased efficacy (27).

A similar analysis of the RhlR LBP revealed that the receptor uses a distinct set of mechanisms to select ligands (10). Indicative of this, RhlR binds to C_4_- and C_6_HSL with low affinity; the C_4_HSL-RhlR interaction has an EC_50_ value of ∼5 μM compared to an EC_50_ value of ∼2 nM for 3OC_12_HSL-LasR, indicating that the different LBPs select ligands via different biochemical constraints (30). Indeed, when RhlR variants were expressed that were expected to enhance ligand binding, as gleaned from LasR ligand selection mechanisms, RhlR exhibited constitutive behavior. RhlR Y64F/W68F/V133F was fully active in a ligand-independent manner, indicating an LBP distinct from other characterized members of the LuxR-type family of receptors. Subsequent biochemical analyses revealed that RhlR could be stably expressed in the presence of the synthetic agonist meta-bromo thiolactone (mBTL) (10,11). The increased stability of the RhlR:mBTL complex is likely the result of enhanced hydrophobic contacts that are absent in the RhlR LBP when it is complexed with native AHL ligands. The purification of RhlR:mBTL was an important step in characterizing other aspects of RhlR function, facilitating the discovery that RhlR binds to the metallo-D-hydrolase PqsE, a known regulator of RhlR function, to enhance the affinity of RhlR for promoter DNA (10,31–36). Full transcriptional activation of RhlR requires ligand binding and PqsE binding in both *P. aeruginosa* and heterologous *E. coli* reporter systems (34,35,37).

Previous structural analyses of RhlR focused on the RhlR-PqsE interface and the role of PqsE in activating RhlR at certain promoters (33,36). In this study, we aim to understand the mechanism of RhlR ligand selection outside of the well-characterized residues (Y64/W68/V133) (Figure 1C). We discover RhlR variants at position T58, specifically substitutions to valine, leucine, and isoleucine, that result in enhanced sensitivity to C_4_HSL. We use the hypersensitive form of RhlR to better understand RhlR promoter binding and gene transcription activation at the *phzABCDEFG1/2* operons, genes that encode the enzymes responsible for the production of the phenazine pyocyanin. Recent work from our group discovered three important characteristics of phenazine gene regulation and pyocyanin production: 1) elevated levels of C_4_HSL lead to decreased levels of pyocyanin production (38); 2) RhlR can bind at the *phzA1* and *phzA2* promoters in the absence of C_4_HSL or PqsE, but not both (37); and 3) RhlR requires both C_4_HSL and PqsE to activate transcription initiation from the *phzA1* and *phzA2* sites (37). Here, we show that the expression of RhlR variants with increased sensitivity to C_4_HSL results in a decrease of *phz1/2* gene expression. Interestingly, the differential gene expression from these sites resulted in the loss of pyocyanin production, but not other QS-dependent traits. We perform ChIP-seq to establish the C_4_HSL-RhlR binding site hierarchy at these loci and assess the phenotypic consequences of that signaling hierarchy. In total, these data establish the molecular basis for the timely expression of RhlR-C_4_HSL dependent genes and the role for PqsE levels in modulating this expression.

## RESULTS

### RhlR ligand binding pocket mutational analyses reveal AHL hypersensitive variants

Previous analysis of the RhlR LBP focused on highly conserved residues that were known from other AHL-binding proteins to be important contacts for AHL molecules, particularly the lactone head group. We performed conservation analyses on 300 LuxR-type receptors to determine variable and moderately conserved residues in the LBP (Figure 1B) and, in conjunction with our previously determined experimental structure of WT RhlR bound to the synthetic agonist mBTL that was then used for docking C_4_- and C_6_HSL. Based on our docking simulations, C_4_- and C_6_HSL occupy a similar position in the LBP as mBTL, a highly efficacious activator of RhlR (Figure 1C and Figure S2). We targeted residues A44, G46, T58, L69, Y72, D81, I84, W96, W108 (Figure 1B and 1C). These residues were selected because of their proximity to the acyl-tail of C_4_HSL as well as the differences in conservation that are observed at these sites. We used an *E. coli* luciferase reporter assay to monitor ligand-dependent activation of WT RhlR and RhlR variants. WT RhlR and RhlR variants were expressed under the control of an arabinose-inducible promoter. Transcription was monitored by light production via the expression of the *luxCDABE* operon controlled by the RhlR-dependent promoter of *rhlA* (Figure S1). Binding of RhlR to the *rhlA* promoter is highly dependent on the C_4_HSL-RhlR interaction, based on ChIP-seq analyses performed in *P. aeruginosa* as well as previously published *E. coli* reporter systems (34,39). After expressing WT RhlR and RhlR variants in the *E. coli* reporter system, we performed 10-point dose response assays using C_4_HSL to calculate the concentration of C_4_HSL required to activate the reporter system to 50% maximum levels (EC_50_ value) (Table 1). As expected, transcription activation by WT RhlR increased with increased C_4_HSL levels with an EC_50_ value of 9 μM. We observed a range of responses; Figure 1C depicts the targeted residues, which are colored based on the effect of amino acid substitution. Here, we focus on the substitutions that resulted in a decrease in the EC_50_ value (*i.e.*, enhanced reporter activation was observed; pink sticks in Figure 1C) or that resulted in an EC_50_ that could not be calculated with the concentrations that were tested (*i.e.*, no reporter activation was observed; orange sticks in Figure 1C).

**Table 1.** EC_50_ values for RhlR and RhlR variants +/- PqsE.

The range of responses to substitutions are best highlighted by changes at residue T58 (Figure 2A). RhlR T58 variants with substitutions to residues with the medium-sized hydrophobic side chains valine, leucine, and isoleucine resulted in decreased EC_50_ values compared to WT RhlR, indicating an enhanced binding to C_4_HSL that resulted in the increased activation of RhlR (Figure 2A; Table 1). The RhlR T58V variant was particularly intriguing because valine occupies the same volume in the LBP but lacks the hydrophilic side chain of threonine. Thus, there is some level of hydrophobic interactions that can be tolerated at this site. Interestingly, increasing the hydrophobicity at this site by increasing the side chain length by one or two carbons (leucine and isoleucine, respectively) at T58 further decreased the EC_50_ value of RhlR (Table 1). However, there was a limit to the additive effect a hydrophobic side chain could have on RhlR activation; RhlR T58F had a slightly increased EC_50_ value (Figure 2A; Table 1). Conversely, the loss of hydrophobicity at the site was not tolerated; a T58G substitution resulted in a complete loss in C_4_HSL activation of RhlR (Figure 2A). Furthermore, enhancing the hydrophobicity of side chains in this particular subregion of the LBP did not generally correlate with enhanced activation by C_4_HSL as RhlR A44M, RhlR A44W, RhlR G46W, and RhlR G46M were not activated by C_4_HSL in the reporter system even at the highest dose of C_4_HSL that was tested (Figure 2A; Figure S3; Table 1). We expect that large hydrophobic residues at A44 are not tolerated, as they prevent stable interactions with the acyl tail of AHLs, even that of C_4_HSL. Going forward, we use the RhlR A44M variant as a control for a RhlR variant that cannot be activated by C_4_HSL. Thus, the accessible volume of the LBP surrounding T58 appears to drive the lower efficacy of C_4_HSL activation of RhlR compared to other LuxR-type receptor/AHL pairs (28,30,40,41). Indeed, of the LuxR-AHL experimentally determined structures this region of the LBP is typically occupied by the acyl tail of long-chain AHLs or a hydrophobic residue (5,28,29,33,42,43), a point we return to in the discussion.

**Figure 2.**
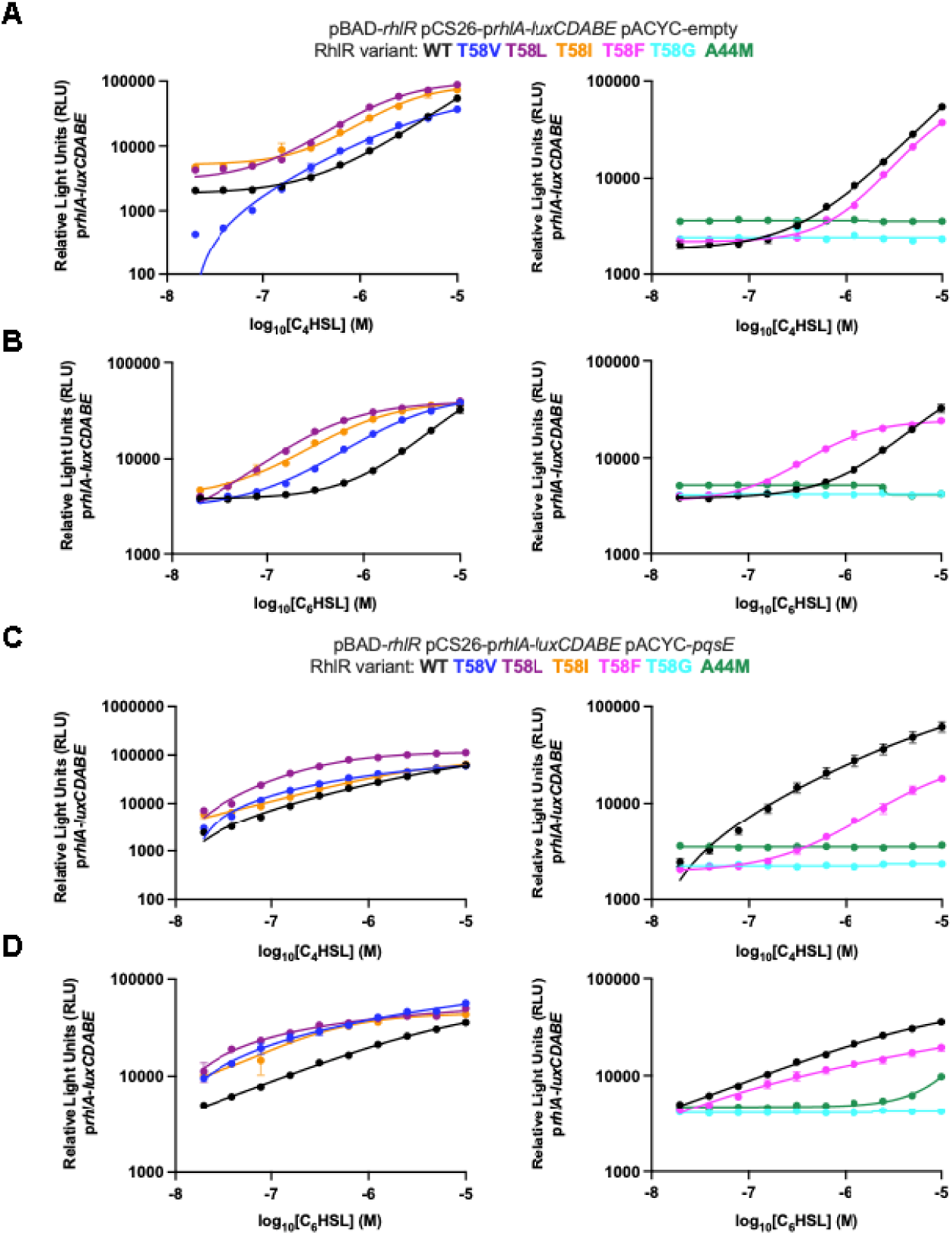
Mutational analysis of the RhlR LBP reveals AHL hypo- and hypersensitive variants. RhlR-controlled bioluminescence was measured in *E. coli*. Arabinose-inducible RhlR was expressed from one plasmid and a p*rhlA*-*luxCDABE* reporter construct was carried on a second plasmid to monitor transcriptional activity. 0.1% arabinose was used to induce RhlR. **A)** RhlR-dependent bioluminescence was measured for WT RhlR (black) and RhlR variants (T58V = blue, T58L = purple, T58I = orange, T58F = magenta, T58G = cyan, A44M = green) in response to increasing concentrations (µM) of C_4_HSL. **B)** RhlR-dependent bioluminescence was measured for WT RhlR (black) and RhlR variants (T58V = blue, T58L = purple, T58I = orange, T58F = magenta, T58G = cyan, A44M = green) in response to increasing concentrations (µM) of C_6_HSL. **C)** RhlR-dependent bioluminescence was measured for WT RhlR (black) and RhlR variants (T58V = blue, T58L = purple, T58I = orange, T58F = magenta, T58G = cyan, A44M = green) in response to increasing concentrations (µM) of C_4_HSL and the presence of overexpressed PqsE from the pACYC plasmid. **D)** RhlR-dependent bioluminescence was measured for WT RhlR (black) and RhlR variants (T58V = blue, T58L = purple, T58I = orange, T58F = magenta, T58G = cyan, A44M = green) in response to increasing concentrations (µM) of C_6_HSL and the presence of overexpressed PqsE from the pACYC plasmid.

To determine the role of hydrophobicity in the LBP adjacent to T58, we assessed the orientation of C_6_HSL in the RhlR binding pocket using a docking simulation. We note that C_6_HSL is a naturally occurring AHL in *P. aeruginosa*, produced by the RhlI synthase that synthesizes C_4_HSL, albeit at significantly lower levels (38). It was previously shown that C_6_HSL can activate RhlR in *E. coli* reporter systems (44). Consistent with that, C_6_HSL activated RhlR in our reporter system with an EC_50_ value 5.5-fold lower than C_4_HSL (Figure 2B; Table 1). We note that while the EC_50_ value for C_6_HSL was lower than C_4_HSL, C_6_HSL did not induce maximal activation of RhlR in the reporter system compared to C_4_HSL (*i.e.*, lower concentrations of C_6_HSL induced higher levels of RhlR-dependent gene expression in the reporter system than the equivalent concentration of C_4_HSL, but the top concentration of C_4_HSL induced more RhlR-dependent gene expression than the top concentration of C_6_HSL). We hypothesize that the activation of RhlR by high levels of an AHL could repress its function as a transcriptional activator by a yet unknown mechanism, such as oligomerization at the promoter binding site (45), which we are currently pursuing but is beyond the immediate scope of this study. Consistent with increased hydrophobic content driving receptor-ligand interactions in the RhlR LBP, C_6_HSL activated RhlR T58V/I/L variants with higher efficacy than C_4_HSL (Figure 2B; Table 1). Interestingly, RhlR T58F had a lower EC_50_ value with C_6_HSL compared to C_4_HSL (Figure 2B; Table 1); again, with the caveat that induction was higher at lower concentrations of C_6_HSL, but C_4_HSL induced maximal light production at the top concentration. Together, these results indicate that the LBP of RhlR can be manipulated to enhance or reduce AHL-dependent activation. Below, we focus on RhlR residues T58 and A44 because they are representative of AHL hyper- and hypo-sensitive RhlR variants. Furthermore, RhlR T58 and A44 variants are biochemically tractable, making them amenable to *in vivo* and *in vitro* assessments of function.

#### The PqsE-RhlR interaction enhances transcriptional activity of some RhlR variants but not others

RhlR requires the accessory binding protein, PqsE, for maximal activity. To determine if PqsE expression altered the activation of RhlR variants, we assessed AHL-dependent RhlR activation of the p*rhlA*-*luxCDABE* reporter in the presence of a third plasmid that constitutively expresses PqsE in *E. coli*. In this assay, PqsE enhances C_4_HSL-RhlR-dependent *rhlA* reporter expression by approximately 2-fold (39). Indeed, the RhlR variants T58V/L/I that displayed enhanced activation by C_4_- and C_6_HSL (Figures 2A and 2B) relative to WT RhlR could be further enhanced by the expression of PqsE (Figures 2C and 2D). RhlR variants that were still active but with increased EC_50_ values for C_4_HSL could have activity enhanced by PqsE. RhlR variants at residue L69 best exemplify this trend, as RhlR L69D/K/M initiated varying levels of transcriptional activity when treated with C_4_HSL and all variants were similarly enhanced by the expression of PqsE (Figure S4). Interestingly, RhlR T58F was not enhanced to the same extent by the expression of PqsE (Figure 2C, 2D). Thus, alterations to the LBP can diminish the additive effect of PqsE-dependent transcription, indicating a potential role of allosteric conformational changes dependent on ligand occupancy. Additionally, the T58G and A44M RhlR variants that could not activate transcription of the reporter did not have function restored by the expression of PqsE (Figures 2C and 2D). A similar trend was observed for other RhlR variants that resulted in a loss of activation by C_4_HSL; RhlR variants G46W, Y72A/T, D81K/N, and W96A (Figure 1C; orange) were recalcitrant to C_4_HSL at all concentrations tested, and activation could not be restored by PqsE (Figures S3 and S5). RhlR variants at residues I84 and W108 could not be activated by C_4_HSL and, thus, light levels remained low at all concentrations that were tested. Unlike the aforementioned variants, RhlR I84M/W and RhlR W108F/Y were activated by the expression of PqsE (Figure S4). Together, these results indicate that different RhlR LBP variants are defective in activating the *rhlA* reporter, likely for different reasons, especially as it relates to activation by PqsE, a point we address with biochemical analyses.

#### Purification of WT RhlR and RhlR variants bound to C_6_HSL reveals connection between PqsE- and AHL-binding

To determine why certain inactive RhlR variants could not be fully rescued by the expression of PqsE in the *E. coli* reporter system, we assessed WT RhlR and RhlR variant solubility in the presence of AHL. Previous attempts to solubilize RhlR in the presence of C_4_HSL were unsuccessful. Given that C_6_HSL can activate RhlR, we performed a RhlR solubility test on RhlR expressed in *E. coli* with C_6_HSL added at the time of induction, a key step to purifying other LuxR-type proteins such as LasR with 3OC_12_HSL. RhlR was soluble in the presence of C_6_HSL (Figure 3A). We hypothesized that the RhlR variants that could not be fully rescued by PqsE were either insoluble or incapable of interacting with PqsE. To test the former, we expressed RhlR variants in the presence of excess C_6_HSL and assessed protein solubility. Many of the RhlR variants, including T58V, were as soluble as WT RhlR. The levels of RhlR variants A44M, L69K, I84M, and W108Y in the soluble fraction were significantly reduced compared to WT RhlR (Figure 3A). Thus, we hypothesize that in the *E. coli* reporter system lacking PqsE, RhlR A44M, L69K, I84M, and W108Y were not binding to AHL and/or were insoluble, resulting in the loss of reporter activity (Figure S4). We next assayed the ability of PqsE to stabilize RhlR variants that were bound to C_6_HSL. We expressed *pqsE* and *rhlR* from the pETDuet-1 vector. This more accurately simulates the *E. coli* reporter system containing *pqsE* than the results from Figure 3A because *pqsE* and *rhlR* are co-expressed under the control of the same T7 promoter elements to ensure roughly equivalent protein levels. All RhlR variants were expressed and soluble, comparable to WT RhlR in the presence of C_6_HSL and PqsE, indicating that PqsE can have a stabilizing effect on RhlR (Figure 3B). However, despite this stability relative to WT RhlR, PqsE could not fully rescue these RhlR variants to WT RhlR levels in the *E. coli* reporter assay (Figure S4).

**Figure 3.**
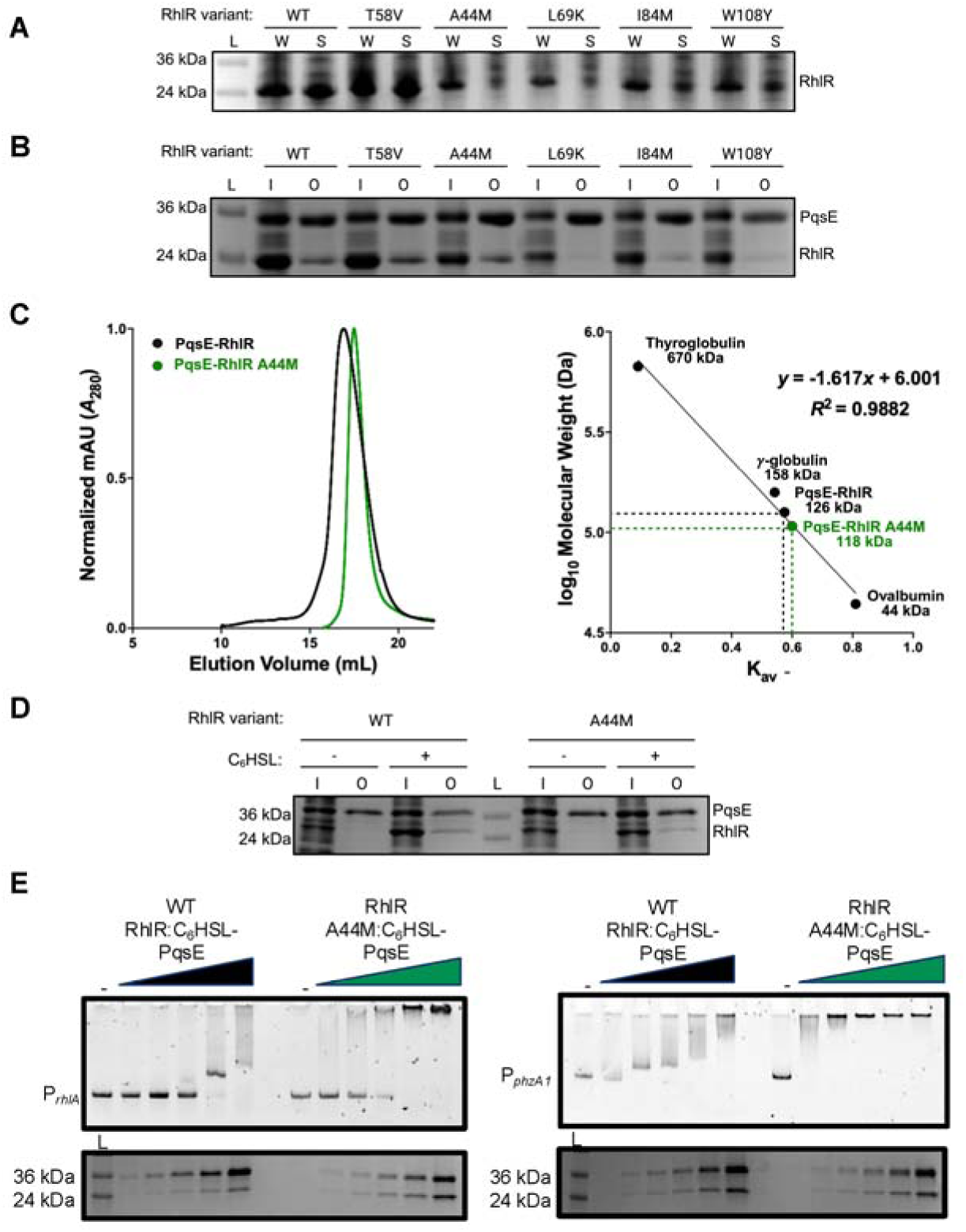
Purification and biochemical characterization of AHL hypo- and hypersensitive RhlR variants. **A)** Total protein comparison of whole cell lysate (W) and soluble (S) fractions from *E. coli* cells overexpressing WT RhlR and RhlR variants in the presence of 200 μM C_6_HSL during induction and lysis. “L” denotes ladder and the 26 and 34 kDa bands are designated. **B)** SDS-PAGE of cell lysates before (input = “I”) and after (output = “O”) affinity purification on Ni-NTA resin. Shown are WT 6x-His-PqsE-containing lysates, which were combined with lysates containing WT RhlR or RhlR variants bound by C_6_HSL. In all affinity purification experiments, RhlR does not carry a tag. “L” denotes ladder and the 26 and 34 kDa bands are designated. **C)** (left) SEC analysis of WT RhlR (black) and RhlR A44M (green) proteins using a Superose-6 column, with protein elution volumes measured by absorbance at 280 nM (*A*_280_, *y*-axis) as a function of retention volume (mL, *x*-axis). Chromatogram traces are representative of three independent purifications for WT RhlR and RhlR A44M. Traces were normalized to a maximum value of 1 for clarity. (right) Molecular weight calculations for WT RhlR (black) and RhlR A44M (green) based on molecular weight standards run using a Superose-6 column. **D)** Same as **C)** but no C_4_HSL was added at the time of induction. **E)** EMSA analysis of the *rhlA* (left) or *phzA1* (right) promoter DNA alone (minus symbol, left lane) with increasing concentrations of purified WT RhlR:C_6_HSL bound to PqsE (black) or RhlR A44M:C_6_HSL bound to PqsE (green). A representative SDS-PAGE gel shows protein levels for each sample (bottom). “L” denotes ladder and the 26 and 34 kDa bands are designated. Final concentrations of WT RhlR and RhlR A44M were: 0.094 , 0.187, 0.375, 0.75, 1.5, and 3 µM per reaction.

To determine if RhlR variants with substitutions in the LBP could still interact with PqsE, we used the pETDuet co-expression system to express RhlR:C_6_HSL and PqsE and then performed an affinity chromatography pulldown experiment. All of the amino acid substitutions described here are buried in the LBP and do not make direct contact with PqsE. As expected, WT RhlR and RhlR T58V interacted with PqsE at equivalent levels (Figure 3B). Surprisingly, RhlR A44M interacted with PqsE similar to WT RhlR and PqsE (Figure 3B). Thus, RhlR A44M was soluble and capable of binding PqsE but not being activated by AHL, indicating that binding to AHL likely induces an allosteric effect on RhlR to drive transcription. Conversely, the RhlR variants L69K and W108Y were significantly diminished in binding to PqsE, which is consistent with PqsE being incapable of fully rescuing RhlR variant activity in the *E. coli* reporter system (Figure 3B and Figure S4). Thus, PqsE is capable of stabilizing RhlR and enhancing its activity as a transcriptional activator in cases when an amino acid substitution in the LBP renders RhlR defective in AHL-dependent activation. Lastly, the RhlR variant I84M had a modest defect in PqsE binding, but this was sufficient to rescue RhlR variant activity in the *E. coli* reporter system (Figure 3B, Figure S4). In total, these results indicate that AHL and PqsE exert different effects on RhlR activation that can be de-coupled. However, the outlier, RhlR A44M, which can bind to PqsE but not be activated by C_6_HSL, indicates that AHL binding is required to drive transcription. This is consistent with our previous ChIP-seq results that identified the *rhlA* promoter as being strongly dependent on AHL for RhlR binding and gene expression (37).

To further characterize the RhlR A44M variant, we purified WT RhlR and RhlR A44M bound to C_6_HSL. Previous purifications of WT RhlR relied on the presence of a highly efficacious AI analog to obtain soluble and stable RhlR. Indeed, this analog was required to determine the RhlR structure (33). However, we recently solubilized RhlR bound to C_6_HSL by adding AHL at the time of induction, which is standard when purifying LuxR-type receptors, as well as during cell lysis and solubilization (46). We used this approach to purify WT RhlR with high purity and yields (Figure 3C and Figure S6). Additionally, we purified RhlR A44M bound to PqsE (Figure 3C; Figure S6). Interestingly, C_6_HSL was required during induction and cell lysis to yield stable RhlR A44M bound to PqsE (Figure 3D), indicating that RhlR A44M could bind to C_6_HSL but could not be activated by AI binding. To further characterize RhlR A44M function, we performed electrophoretic mobility shift assays (EMSA) to determine the DNA binding capability of both WT RhlR and RhlR A44M bound to PqsE and C_6_HSL. WT RhlR:C_6_HSL-PqsE complex bound to both the *rhlA* and *phzA1* promoters (Figure 3E), as previously described. Interestingly, RhlR A44M and PqsE induced a super shift of both the *rhlA* and *phzA1* promoters, likely reflective of enhanced binding for the Rhl box motif sequence or enhanced ability to oligomerize on the DNA. We address the ability of RhlR A44M to bind DNA in our ChIP-seq analyses below. These biochemical experiments represent two important advances to understanding RhlR structure-function: 1) WT RhlR can be purified with a “native” AI in the presence of PqsE and 2) a variant of RhlR incapable of being activated by AI can still be soluble in the presence of the AI and PqsE and, thus, is purifiable.

#### AHL hypersensitive RhlR variants display altered phenotypic traits in P. aeruginosa

To assess the consequences of RhlR hypersensitivity to AHL molecules, we performed phenotypic analyses of well-characterized RhlR-dependent QS traits, such as pyocyanin, biofilms, swarming, and rhamnolipid production, in strains expressing *rhlR* mutants from the native *rhlR* locus. The RhlR variant proteins were expressed to similar levels as WT RhlR (Figure S7A). We previously showed that pyocyanin production via the regulation of the *phzABCDEFG1/2* operon by RhlR was sensitive to levels of C_4_HSL; C_4_HSL concentrations below ∼500 nM and above ∼5 µM resulted in a loss of pyocyanin production (38). Thus, we hypothesized that expression of a RhlR variant with higher affinity for C_4_HSL would repress pyocyanin production at high-cell density due to hyperactivation of the Rhl system. Indeed, we observed this phenomenon in strains producing their own C_4_HSL (*i.e*., an otherwise WT background), as strains expressing RhlR T58L but not RhlR T58V, produced less pyocyanin than WT (Figure 4A). Additionally, a strain expressing RhlR A44M was unable to produce detectable levels of pyocyanin, indicating a loss of RhlR-dependent signaling via C_4_HSL (Figure 4A). To ensure that the increased sensitivity to C_4_HSL by the RhlR variants and not the dysregulation of *rhlI*, which would alter the C_4_HSL concentrations in these backgrounds, led to the decrease in pyocyanin production, we measured pyocyanin levels in a Δ*rhlI* strain expressing WT RhlR and RhlR variants in the presence of 2 µM C_4_HSL. Supplementation of a Δ*rhlI* strain expressing RhlR T58V or T58L with 2 µM C_4_HSL resulted in less pyocyanin produced by the culture compared to a Δ*rhlI* strain expressing WT RhlR treated with 2 µM C_4_HSL (Figure 4B), indicating that RhlR sensitivity to C_4_HSL is important for the regulation of phenazine gene expression. Thus, pyocyanin production can be suppressed when RhlR is overactivated by AHL.

**Figure 4.**
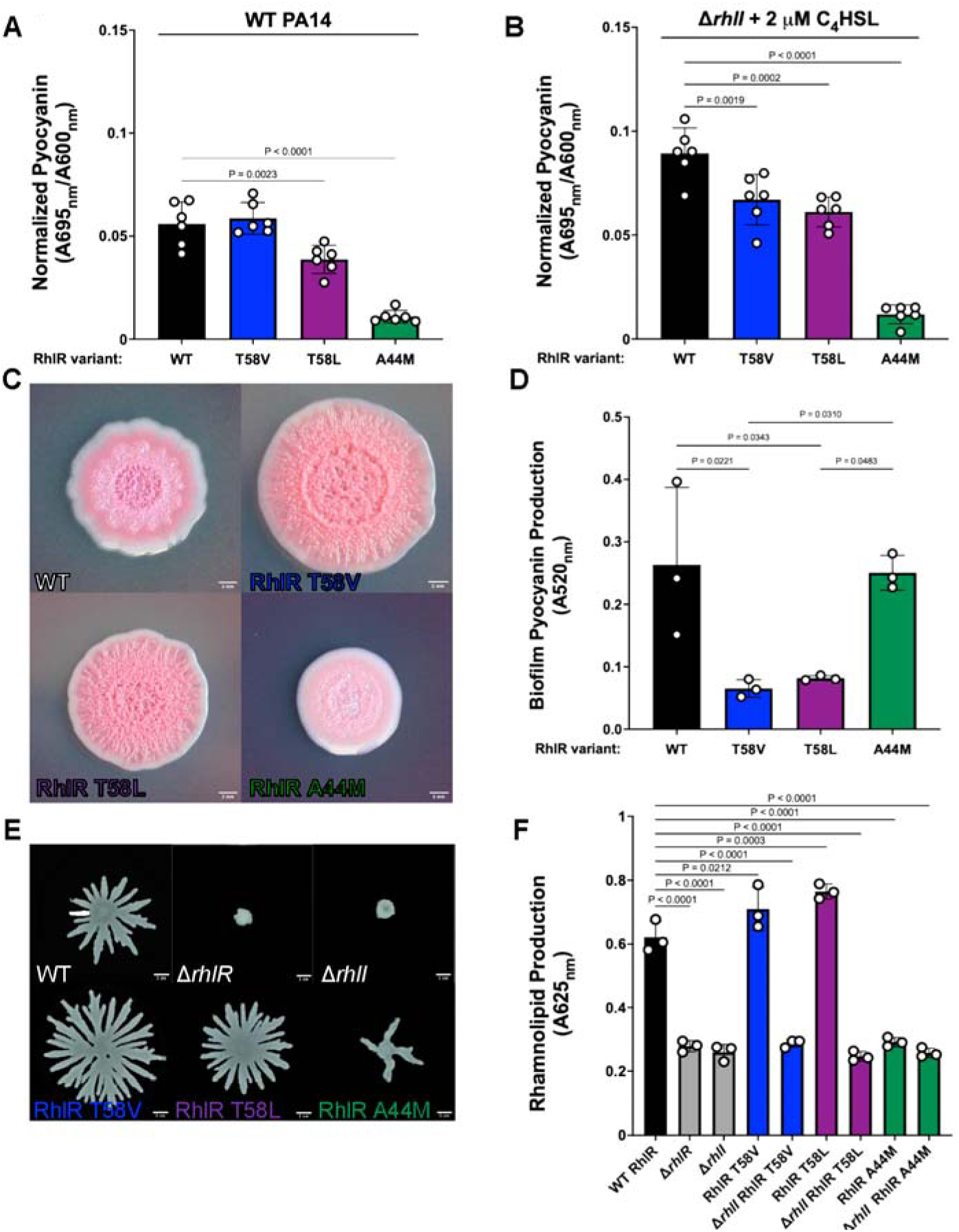
Phenotypic analysis of AHL hypo- and hypersensitive RhlR variants in *P. aeruginosa*. **A)** Pyocyanin production was measured in WT *P. aeruginosa* strains expressing WT RhlR or the designated RhlR variants. Pyocyanin was quantified as pigment production (OD_695 nm_) divided by cell density (OD_600 nm_). **B)** Same as A) except pyocyanin production was measured in a *P. aeruginosa* Δ*rhlI* strain expressing WT RhlR or the designated RhlR variants in the presence of 2 μM C_4_HSL. **C)** Representative images of Congo red colony biofilm plates for strains of *P. aeruginosa* expressing WT RhlR, RhlR T58V, RhlR T58L, and RhlR A44M. **D)** Pyocyanin production from extractions obtained from colony biofilms grown in **C**. Extracted pyocyanin was measured at 520_nm_. **D)** Rhamnolipid production of the supernatants from the designated strains grown in LB for 24 hours using Victoria Pure Blue BO and measuring absorbance of the dye at OD_625 nm_. **E)** Swarming motility assay on 0.4% agar using *P. aeruginosa* strains expressing WT RhlR or the designated RhlR variants. Statistical analyses for all assays were performed using an ordinary one-way ANOVA with a Dunnett’s multiple comparisons test. All comparisons were made to the WT control strain; comparisons that were deemed not significant by these analyses are not shown.

To understand if C_4_HSL sensitivity affected phenazine production in the context of biofilms, we performed analyses on the above strains to assess their ability to produce pyocyanin and form colony biofilms on Congo red plates. WT *P. aeruginosa* produced a stereotypical colony biofilm with a rugose center and smooth outer edge (Figure 4C). WT *P. aeruginosa* strains expressing RhlR T58V and T58L variants produced hyper-rugose colony biofilms with a smaller smooth periphery (Figure 4C). Conversely, a WT *P. aeruginosa* strain expressing the RhlR A44M variant produced markedly different colony biofilms with a largely smooth texture (Figure 4C). The colony biofilm formed by the strain expressing RhlR A44M further confirms that RhlR signaling is intact via PqsE but not through the activation by C_4_HSL. Furthermore, it was previously shown that a Δ*pqsE* strain phenocopies the hyper-rugosity of a Δ*rhlR* strain. Thus, we hypothesize that the strain expressing a RhlR T58L variant is hyper-rugose because its enhanced binding to C_4_HSL has a subsequent effect on PqsE levels, which we address below. Additionally, these phenotypes are consistent with previous findings that showed that Δ*rhlI* strains exhibit a smooth colony morphology, respectively.

Colony morphology and rugosity often correlate with phenazine production; deletion of the phenazine operons leads to a hyper-rugose colony morphology similar to that of a Δ*rhlR* strain (47). The hyper-rugose phenotype is thought to function as a mechanism to compensate for the loss of pyocyanin, which can act as a terminal electron acceptor under anoxic conditions. Thus, we hypothesized that strains expressing RhlR variants that were hyper-sensitive to C_4_HSL would produce less pyocyanin compared to WT or the strain expressing the RhlR A44M variant (Figure 4D). We extracted pyocyanin from the Congo red biofilm plates and measured the absorbance at 520_nm_. Strains expressing the RhlR T58V and T58L variants produced significantly less pyocyanin than the WT strain or the strain expressing the RhlR A44M variant (Figure 4D; Figure S7B). Thus, while the strain expressing the A44M variant is defective in pyocyanin production in planktonic culture, it is still capable of producing pyocyanin when grown in a biofilm. This is consistent with previous findings that showed that *rhlI* mutants are smooth, express the phenazine genes, and produce pyocyanin, which is in contrast to *rhlR* mutants grown as planktonic or biofilm cultures (10,47). Thus, the strain expressing the RhlR A44M variant can still drive gene expression under certain environmental conditions but, given that colony morphology is not the same as the WT phenotype, it is likely context specific (*i.e.*, only certain RhlR-dependent genes can be regulated). This establishes that colony biofilm formation and pyocyanin production correlate with RhlR sensitivity to AHL.

To determine if sensitivity to C_4_HSL specifically repressed pyocyanin production or if it was a generalized response to suppress other QS traits, we performed additional phenotypic analyses of the above strains to assess the ability of *P. aeruginosa* to swarm on low percentage agar plates, a trait known to be dependent on the RhlR-AHL interaction (48,49). As expected, WT *P. aeruginosa* was capable of swarming and a Δ*rhlI* strain was not (Figure 4E; Figure S7C). Distinct from the observed pyocyanin production phenotypes, strains expressing RhlR T58V and T58L were capable of swarming and matched the total swarming diameter of WT *P. aeruginosa* (Figure 4E; Figure S7C), indicating that not all *P. aeruginosa* RhlR-dependent QS phenotypes are suppressed by hypersensitivity to AHL. Conversely, a strain expressing RhlR A44M was incapable of swarming to WT levels, consistent with the inability of the variant to be activated by AHL (Figure 4E). We next correlated the ability to swarm with rhamnolipid production, a well-characterized C_4_HSL-RhlR-dependent trait. The ability of strains to swarm was consistent with rhamnolipid production as measured by the absorbance of Victoria Blue PO with cell-free supernatant (Figure 4F). Furthermore, the hyperactive strains required the presence of C_4_HSL to produce rhamnolipids, as all RhlR variants in a Δ*rhlI* background were defective in rhamnolipid production. Thus, rhamnolipid production and the subsequent swarming of the colony correlates with RhlR sensitivity to AHL.

#### Enhanced selectivity of RhlR for C_4_HSL leads to differential gene expression

To determine the effects of hypo- and hypersensitivity to C_4_HSL on RhlR-dependent transcription activation in *P. aeruginosa*, we performed RNA-seq on Δ*rhlI* strains expressing WT RhlR, RhlR T58L, and RhlR A44M in the presence of 2 µM C_4_HSL. We chose this approach to better control the levels of C_4_HSL across samples. All gene expression analyses were done relative to a Δ*rhlI* strain without exogenous supplementation of AHL. Thus, we examined the total increase in C_4_HSL-dependent gene regulation. We observed increased *rhlA* RNA levels in strains expressing WT RhlR and the RhlR T58L variant compared to Δ*rhlI* without C_4_HSL or a strain expressing the RhlR A44M variant (Table 2, Table S1). Additionally, many other genes that were previously reported as dependent on the RhlI-synthesized AI showed similar trends; *lasB*, *mexG*, *katAB,* and *chiC* were expressed at higher levels when a hypersensitive RhlR variant was expressed (Figure 5A; Table 2). To corroborate our RNA-seq data, we built a *rhlA* promoter fusion to mScarlet and assayed reporter levels in the same genetic backgrounds as our transcriptomic experiment (Figure S7D). Strains expressing hypersensitive RhlR variants produced similar levels of fluorescent signal relative to the WT strain, while a strain expressing a hyposensitive RhlR variant had diminished levels of fluorescent reporter (Figure S7D). Furthermore, the *rhlA* transcript levels in both of our assays are consistent with the swarming and rhamnolipid production data in Figure 4E and 4F. Additionally, we note that not all genes that were upregulated in a C_4_HSL-dependent manner are the result of direct regulation by RhlR, as many genes with altered expression did not have a corresponding RhlR binding site in their promoter region, as defined by our previous ChIP-seq analyses (37).

**Figure 5.**
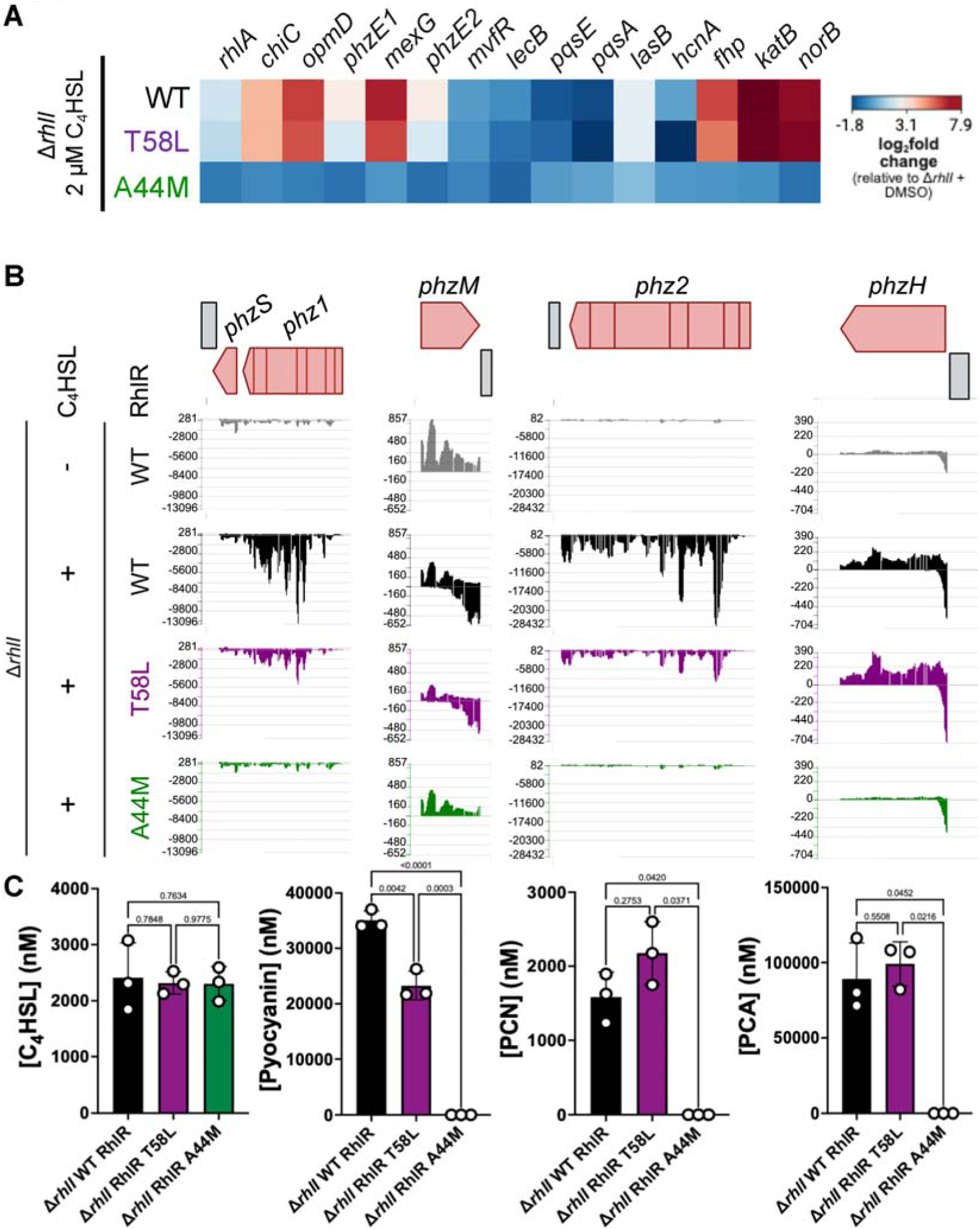
RhlR variant activation alters expression of phenazine genes and pyocyanin production. **A)** Heat map of normalized gene expression between Δ*rhlI* strains expressing WT RhlR in the presence of 2 μM C_4_HSL with Δ*rhlI* strains expressing RhlR A44M or RhlR T58L variants in the presence of 2 μM C4HSL. **B)** Signal map profiles of the designated strains showing the level of mapped RNA reads to all phenazine genes: *phzABCDEFG1/2* operons*, phzM*, *phzS*, and *phzH*. **C)** Absolute concentrations of C_4_HSL, pyocyanin, PCA, and PCN from cell-free supernatants of WT strains expressing WT RhlR and RhlR variants T58L and A44m using ultra-high-performance liquid chromatography (UHPLC) coupled with a high-resolution mass spectrometer (HRMS). All strains were supplemented with 2 μM C_4_HSL. Bars represent the mean of three biological replicates.

**Table 2.** Relative gene expression for QS genes of interest.

Many of the *phz* genes were exceptions to the above trend of WT or higher levels of gene expression in backgrounds that were hypersensitive to C_4_HSL. The enzymes responsible for the synthesis of pyocyanin are encoded by the *phzABCDEFG1/2* operons, *phzH*, *phzM*, and *phzS*. We previously mapped RhlI-dependent RhlR binding sites upstream of the *phzA1*, *phzA2*, and *phzH* genes and within the *phzB1* and *phzB2* coding sequences (37). Thus, we hypothesized that these genes would be sensitive to the addition of C_4_HSL. Consistent with the above phenotypic analyses of pyocyanin production, *phz* gene expression was lower in strains of *P. aeruginosa* that expressed a hypersensitive RhlR variant. Notably, the expression of *phzA/B/C1* were especially sensitive, as gene expression levels were comparable to that of a Δ*rhlI* strain without exogenous C_4_HSL (Figure 5B and Figure S7E). A strain expressing T58L had lower *phz* gene expression across all of the *phz* genes with the exception of *phzH*, which encodes the enzyme responsible for converting phenazine-1-carboxylic acid (PCA) to phenazine-1-carboxamide (PCN). All of the genes in the *phzABCDEFG1/2* operons were expressed at least 25% lower in a strain expressing the RhlR T58L variant compared to WT RhlR. To confirm our RNA-seq results of decreased phenazine gene expression, we performed chromosomal replacement of WT *rhlR* with our mutant *rhlR* alleles into *phz1*-mScarlet and *phz2*-mScarlet reporter *P. aeruginosa* strains (Figure S7E). Interestingly, a strain expressing RhlR T58L had an insignificant decrease in *phz1*-mScarlet expression, while *phz2*-mScarlet expression decreased by 4.5-fold (Figure S7E). Thus, the *phz2* promoter is likely more sensitive to increased C_4_HSL signaling. Conversely, both *phz* reporters displayed similar 2-fold decreases in a strain expressing RhlR A44M (Figure S7E). We next measured *phz2*-mScarlet expression in the context of colony biofilms (Figure S8A). These data were consistent with our observations of biofilm morphology and biofilm pyocyanin production (Figures 4C and 4D); *phz2* reporter expression was maximal in the WT strain, significantly decreased in a strain expressing RhlR T58L, and moderately decreased in a strain expressing RhlR A44M (Figure S8B). In total, these data show that RhlR sensitivity to C_4_HSL can be promoter specific and dependent on the environment. We note that *phzH*, *phzA1*, and *phzA2* have separate RhlR binding sites in their respective promoter regions. Thus, RhlR sensitivity to C_4_HSL controls differential *phz* gene expression from the different promoters that contain RhlR binding sites; hyperactivation of RhlR leads to lower levels of *phz* gene expression from the *phzA1* and *phzA2* promoters but not *phzH*.

To determine the relative levels of different phenazines produced as a result of the altered expression of the *phz* genes, we performed liquid chromatography-mass spectrometry (LC-MS) to quantify pyocyanin, PCA, and PCN from planktonic cultures (Figure 5C, Figure S9). Consistent with our spectrophotometric data, pyocyanin levels were lower in a strain expressing RhlR T58L relative to WT RhlR. Given that *phzH* gene expression levels were higher in a strain expressing RhlR T58L compared to WT RhlR, we hypothesized that PCN, the product of PhzH activity, would be elevated in a strain expressing RhlR T58L. Indeed, PCN levels increased by 37% increase in a strain expressing RhlR T58L (Figure 5C). We surmise that the strain expressing RhlR T58L maintains enough gene expression through the *phz1* and *phz2* operons that a sufficient amount of substrate is available for PhzH to synthesize PCN. Thus, we expected that the concentrations of PCA would remain relatively unchanged between WT and RhlR T58L strains and, indeed, WT and RhlR T58L produced nearly identical levels of PCA (Figure 5C). Unlike the strain expressing RhlR T58L, the strain expressing RhlR A44M was deficient in the production of all phenazine metabolites (Figure 5C), which is consistent with the phenotypic and transcriptional data that showed a complete loss of gene expression from all of the *phz* promoters (Figures 4B and 5A). Thus, changes in gene expression can alter the flux of metabolites through the phenazine metabolic machinery, which alters the pool of secreted phenazines, resulting in lower levels of pyocyanin and higher levels of PCN.

#### Decreased levels of PqsE lead to decreased promoter binding for RhlR at phenazine gene promoters

To determine the molecular basis for lower levels of *phz* gene expression in a background that expressed a hypersensitive variant of RhlR, we assessed expression of the *pqsABCDE* operon (50). PqsE is required for optimal expression of the *phz* genes and, thus, pyocyanin production. RhlR was previously characterized as a repressor of the *pqsABCDE* operon. We hypothesized that a C_4_HSL hypersensitive variant would repress *pqsABCDE* expression, which could lead to the observed decrease in *phz* gene expression. Indeed, *pqsA* RNA levels were 20% lower in a strain expressing RhlR T58L compared to WT RhlR (Figure 5A; Table 2). To confirm that the decrease in gene expression led to a decrease in the available PqsE protein that was capable of interacting with RhlR to specifically drive it to the *phz* promoter, we performed western blot analyses to measure PqsE protein levels in strains expressing RhlR variants (Figure 6A). PqsE levels were approximately 2-fold lower in a strain expressing the RhlR T58L variant compared to WT RhlR, indicating that, despite its hypersensitivity to C_4_HSL, RhlR requires a certain amount of PqsE to initiate transcription of the *phz* genes. We recently showed that PqsE dimerization is concentration dependent (46) and that PqsE dimerization is required for complex formation (33). Thus, we hypothesized that when PqsE levels fall below a certain concentration, PqsE homodimerization cannot occur, thereby preventing PqsE-RhlR complex formation and reducing promoter occupancy. To determine if the lower levels of PqsE resulted in less enrichment of RhlR at RhlR-dependent promoters, specifically at the phenazine promoters, we performed ChIP-seq analyses to assess RhlR DNA binding genome wide (Figure 6B, 6C, Figure S10A-C, Table 3). We highlight *phzA1* and *phzM*, which share a promoter, (Figure S10A), *rhlA* (Figure S10B), and *lecB* (Figure S10C) because of the role of the *phz* and *rhlA* promoters in regulating pyocyanin and rhamnolipid production (Figure 4) and because *lecB* is a well-characterized PqsE-dependent promoter (37). As expected, RhlR T58L had lower promoter occupancy than WT RhlR at the *phzA1* promoter, likely due to the lower levels of PqsE present (Figure 6B, S10A). Conversely, RhlR occupancy at the *rhlA* promoter was nearly unchanged from WT RhlR (Figure 6B, S10B), reflecting the ability of RhlR to bind the promoter primarily through its dependency on C_4_HSL. Additionally, RhlR A44M occupancy was more significantly reduced at the *phzA1/phzM* and *rhlA* promoter, consistent with C_4_HSL binding being important for RhlR occupancy at these promoters (Figure 6C, S10A, S10B).

**Figure 6.**
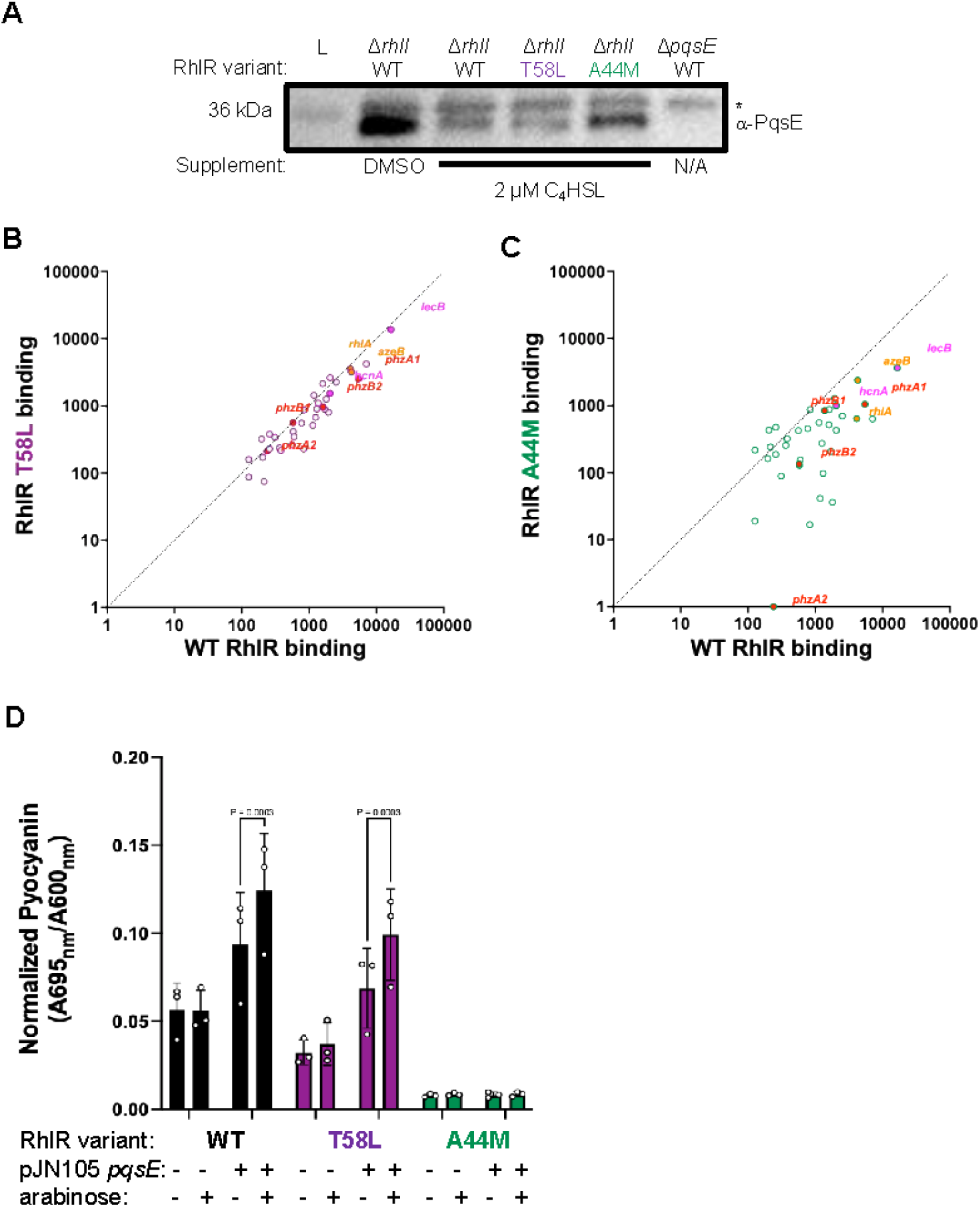
Repression of *pqsE* alters the ability of RhlR to bind to the *phz* promoter. **A)** Western blot analysis depicting PqsE protein levels in the designated strains using a polyclonal antibody raised against purified PqsE. **B)** Comparison of ChIP enrichment from the Δ*rhlI* and Δ*rhlI* RhlR T58L strains or **C)** Δ*rhlI* RhlR A44M, all containing 2 μM of C_4_HSL. **D)** Pyocyanin production was measured in WT *P. aeruginosa* strains expressing WT RhlR or the designated RhlR variants. Pyocyanin was quantified as pigment production (A695 _nm_) divided by cell density (A600 _nm_). Each strain contained a pJN105 empty vector (-) or a pJN105 containing *pqsE* (+) under the control of the pBAD promoter. Arabinose was supplied at a final concentration of 0.1% (v/v) (+) or an equivalent volume of water (-) was supplied to all cultures.

**Table 3.** ChIP-seq analyses for all 40 previously annotated RhlR binding sites.

RhlR T58L did not have significantly altered occupancy at the *lecB* binding site compared to WT, which is counter to our expectations. We hypothesize that reduced PqsE levels would have an effect on RhlR binding to PqsE-dependent promoters (Figure S10C). Furthermore, RhlR occupancy did not correlate with gene expression from these promoters, which we also observed when we previously performed ChIP-seq experiments in *rhlI* and *pqsE* deletion strains. For example, despite RhlR T58L and RhlR A44M occupying the *lecB* promoter at different levels, the expression profiles were nearly identical (Figure S10C, Tables 2 and 3). While it was initially surprising to observe that RhlR T58L could occupy the *lecB* promoter because it is so strongly dependent on the levels of PqsE, we hypothesize that if some number of PqsE dimers exist in complex with RhlR that these complexes would be preferentially driven to the *lecB* promoter. The *lecB* promoter is one of the “preferred” sites for PqsE-RhlR binding in the genome, as determined by map reads to the *lecB* promoter from our previous ChIP-seq data (37). Additionally, we cannot rule out that the increased efficacy of C_4_HSL for RhlR T58L alters RhlR binding site preferences. Future studies aimed at determining the hierarchy of RhlR promoter selection will be needed to confirm this assertion.

To test whether low levels of PqsE were directly leading to the dysfunction in the regulation of the phenazine genes in strains that were hyper-responsive to C_4_HSL, we overexpressed *pqsE* from the pBAD promoter on the pJN105 plasmid in WT, RhlR T58L, and RhlR A44M variant backgrounds of *P. aeruginosa* and assayed for pyocyanin production (Figure 6D). All strains received the equivalent pJN105 empty vector control. As shown in Figure 4, the strain expressing RhlR T58L that contained the empty plasmid produced less pyocyanin than the WT counterpart. We note that we observed an intermediate enhancement of pyocyanin production for the strains expressing WT RhlR and RhlR T58L containing pJN105-*pqsE*, likely reflecting the leaky expression of *pqsE* under the pBAD promoter (Figure 6D). Nevertheless, induction of *pqsE* expression from the pBAD promoter with the addition of arabinose enhanced pyocyanin production in both the WT strain as well as the strain expressing RhlR T58L, as expected, indicating that levels of PqsE are important for driving and finetuning RhlR-dependent transcription (Figure 6D). Lastly, the strain expressing RhlR A44M did not exhibit restored pyocyanin production under any of the conditions tested (Figure 6D), consistent with the results from the *E. coli* reporter system (Figure 2C and 2D) and the phenotypic assays performed in *P. aeruginosa* (Figure 4). Thus, the RhlR A44M variant is unaffected by PqsE levels and represents a completely inactive form of RhlR. In total, RhlR promoter binding must be effectively coupled with RNA polymerase to drive gene expression, and this is likely mediated both by PqsE and allosteric effects induced by C_4_HSL occupancy in the RhlR LBP.

## DISCUSSION

Here, we showed that amino acid substitutions at T58 increase the sensitivity of RhlR to C_4_HSL, which can have significant disparate effects on downstream signaling. We hypothesize that the RhlR LBP reflects an evolutionarily optimized sensitivity to C_4_HSL. It is intriguing that RhlR has such a high EC_50_ value and requires > 1 μM C_4_HSL for QS activation. This is in stark contrast to LasR, a second LuxR-type receptor also found in *P. aeruginosa*. This likely reflects the role of specificity and sensitivity of the two systems. LasR has a high sensitivity for 3OC_12_HSL, but it comes at the cost of decreased selectivity; LasR can be activated by 3OC_8_HSL, 3OC_10_HSL, and 3OC_14_HSL, albeit with reduced sensitivity relative to 3OC_12_HSL (28). We hypothesize that this is due to the placement of LasR in the QS hierarchy; LasR signaling is initiated prior to RhlR activation and as an early sensor for the chemical signaling environment, it might be advantageous to have initial flexibility in AHL binding to robustly initiate QS to outcompete neighboring bacteria that produce non-cognate AHL. Conversely, RhlR sacrifices sensitivity for selectivity and appears to maintain this level of sensitivity under some evolutionary constraint given that we show it could have evolved greater sensitivity. We hypothesize that this is due to the costs associated with producing phenazines (51). More specifically, the overproduction of pyocyanin can be toxic to *P. aeruginosa*. Thus, if RhlR were too sensitive to C_4_HSL and/or overly promiscuous it would be difficult to fine tune or maintain proper regulation of pyocyanin. Additionally, this fine tuning is optimally achieved because RhlR has co-evolved with PqsE to control expression of its regulon. Thus, RhlR coincidence detection via C_4_HSL and PqsE underpins late QS behaviors in *P. aeruginosa* (Figure 7).

**Figure 7.**
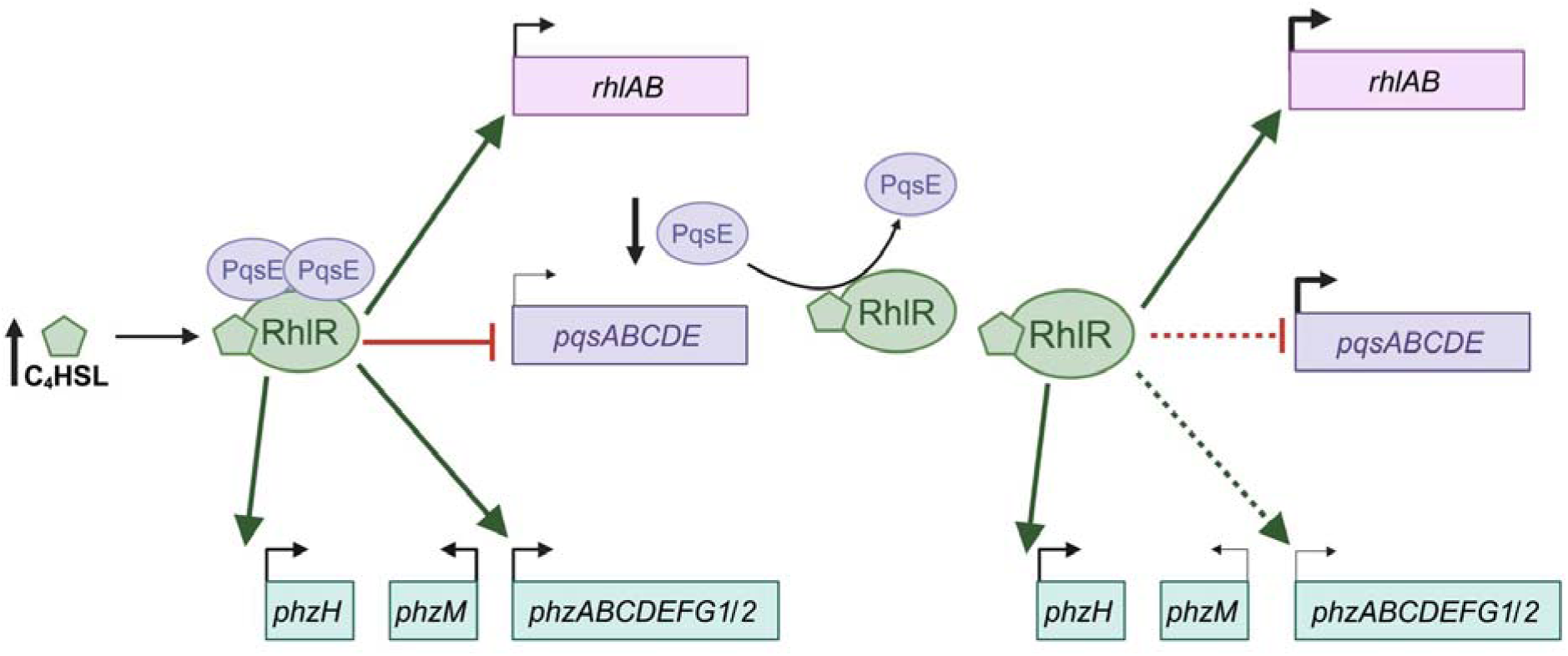
Model for C_4_HSL- and PqsE-dependent gene expression. The levels of C_4_HSL and PqsE tightly regulate the activation of RhlR by altering its binding to specific DNA sites.

We used RhlR hyperactive variants to mimic hyperactivation by C_4_HSL and to determine the crucial role of C_4_HSL sensing in QS progression. The phenazines are subject to complex regulation to allow for adaptation to different environmental conditions. Our data support the unique and dynamic role of RhlR-PqsE regulation; C_4_HSL levels influence both C_4_- and PqsE-dependent activation of different phenazine genes. Specifically, our data showed that an increase in *phzH* expression relative to *phzM* shuttles substrates to the production of PCN. PCN can be reduced and re-oxidated to function as an electron acceptor, thereby helping the cell maintain a balanced redox state (52,53). This is especially important during anaerobic growth and biofilm growth; PCN enables the conversion of glucose and pyruvate to acetate to promote redox homeostasis and ATP synthesis, which is essential for cell survival during fermentative growth (51,54). We hypothesize that C_4_HSL functions as a proxy for this “extreme” environment. During biofilm formation, we expect that C_4_HSL levels would be highest at the center of the biofilm with the physical constraints of the biofilm dampening C_4_HSL diffusion into the greater extracellular milieu. In turn, this would signal to the cells to convert PCA to PCN rather than pyocyanin to ensure the survival of cells grown at the center of the biofilm. Meanwhile, cells at the periphery are actively growing, secreting and sensing freely diffusible C_4_HSL, which subjects this population to the cycling coincidence detection observed in our RhlR-PqsE gene expression data and coincides with the robust production of pyocyanin instead of PCN. Thus, a single coincidence detection system can act as a switch to allow for the formation of different parts of a colony biofilm. Hyperactivation of RhlR by C_4_HSL did not have a similar effect on genes involved in cellular detoxification such as superoxide dismutase, heme oxygenase, nitric oxide reductase, and alkyl hydroperoxide reductase. These are enzymes that would be beneficial to collective cell survival alongside the differential production of phenazines. However, regulation of these genes appears to be indirect, yet through the response of RhlR to C_4_HSL, as the only difference between strains in our RNA-seq experiments was the expression of the hyperactive RhlR variant. Thus, it is intriguing to envision that C_4_HSL levels function as a proxy for the late stages of QS, with survival adaptation to environmental stresses and perturbations being paramount via changes in gene expression, especially in the context of biofilms.

Our chemical-genetic approach revealed important nuances about the RhlR LBP that will be informative for future medicinal chemistry approaches aimed at inhibiting the RhlR-C_4_HSL interaction. In particular, the RhlR A44M variant was insightful because it revealed a portion of the LBP that, when perturbed, could reduce or eliminate the activation of RhlR by C_4_HSL. Importantly, RhlR A44M was soluble in the presence of PqsE and that allowed us to recapitulate the RhlR-PqsE interaction *sans* activating ligand, which had previously been a state of RhlR that could not be purified. In a cellular context, RhlR-PqsE while unliganded was incapable of driving gene expression from any promoter. RhlR A44M could, however, still bind to promoters that were previously shown to be reliant on PqsE for binding. We hypothesize that the RhlR A44M variant induces an allosteric structural change that eliminates the ability of RhlR to engage with RNA polymerase. It was previously postulated that to fully inhibit RhlR function would require suppression of both C_4_HSL and PqsE-dependent traits (1,39). Here, we show that strains expressing RhlR A44M fail to activate C_4_HSL- and PqsE-dependent traits via a single change in the RhlR LBP. Thus, if the pocket surrounding A44 in the RhlR LBP can be targeted with a C_4_HSL-like mimic that occupies the area adjacent to A44 with a hydrophobic moiety, RhlR-dependent gene expression could be effectively abolished to levels comparable to that of a Δ*rhlR*, which would render *P. aeruginosa* non-pathogenic.

## EXPERIMENTAL PROCEDURES

### Phylogenetic tree construction

Two separate lists of protein sequences were retrieved from the National Center for Biotechnology Information (NCBI): one for LuxR receptors and another for their partner LuxI synthases. Each list was aligned using MUSCLE v3.8.1551 and trimmed with trimAl v1.4.1 using default parameters. Maximum-likelihood trees were generated from alignments of 22 sequences for each group, with bootstrap support values calculated with 1,000 replicates using IQ-TREE v1.6.12. Model selection was performed using an automatic substitution model based on the Bayesian information criteria (BIC) score, selecting the LG+G4 model for the LuxR tree and the LG+I+G4 model for the LuxI tree. The trees were visualized and annotated with Interactive Tree of Life (iTOL) v6.9.1.

### Plasmid and strain construction

Plasmids and strains were constructed using standard molecular cloning techniques. Specifically, *rhlR* in the pBAD-A plasmid was mutagenized using overlap extension PCR with site-directed mutagenesis primers designed using the Agilent QuikChange Primer Design Tool. Sequence confirmed pBAD-A-*rhlR* mutants were transformed into an *E. coli* strain already containing pCS26-p*rhlA*-*luxCDABE* and pACYC-*pqsE* or the pACYC empty vector control. We also note that the RhlR and PqsE co-expression construct (pETDuet-1) was cloned to not include RhlR with an N- or C-terminal tag; PqsE was tagged at the N-terminus as it was previously shown to be functional *in vivo*, indicating that it has no effect on the ability of PqsE to bind to RhlR. Table S2 lists the plasmids and strains used in the study. Table S3 lists the primers used in the study.

### *P. aeruginosa* strain construction

Standard cloning and molecular biology techniques were used to generate *P. aeruginosa rhlR* mutations. Introduction of genes encoding RhlR variants onto the *P. aeruginosa* chromosome was achieved using previously published protocols (47). In brief, the entire *rhlR* mutant coding sequence was amplified, digested, and ligated into the pEXG2 vector. The pEXG2 vector containing *rhlR* mutants was then transformed into *E. coli* SM10 λ*pir,* followed by conjugation into the appropriate UCBPP-PA14 *P. aeruginosa* strain background. All strains were confirmed by amplifying the target locus followed by Sanger sequencing. Transcriptional reporters for the *phz1* and *phz2* operons were generated as previously described, with plasmids generously gifted by Dr. Lars Dietrich (55).

### *E. coli* light assay

All *E. coli* strains in this study were grown overnight with shaking in 5 mL LB media supplemented with 100 µg/mL ampicillin, 50 µg/mL kanamycin, and 15 µg/mL tetracycline at 37°C overnight. Overnight cultures were diluted 1:100 into fresh media, grown at 37 °C with shaking for 2 h until reaching OD_600_ = 0.4. Cultures were induced with a final concentration of 0.1% arabinose and 198 μL of the induced culture were mixed with 2 μL of C_4_HSL dilutions in a clear bottom black opaque 96-well plate (Corning) and grown at 37°C with shaking for 4 h. The C_4_HSL dilution series was performed 1:1 in DMSO with a top final concentration of 10 μM. Luminescence and optical density were recorded using a Glomax (Promega) plate reader.

### WT RhlR, RhlR variant, and PqsE protein expression, affinity purification pulldown, and large-scale purification

To determine protein solubility +/- AHL or PqsE, WT RhlR and other RhlR variant genes were cloned into the multiple cloning site 2 of the pETDuet-1 vector using NEBuilder HiFi Assembly cloning kit. To perform the affinity purification pulldown, the 6x-His-*pqsE* gene was cloned using the same cloning method into the multiple cloning site 1 of pETDuet-1 already having either WT *rhlR* or *rhlR* mutant genes in the multiple cloning site 2. Sequence confirmed plasmids were transformed into *E. coli* BL21 (DE3) cells. For both solubility and pulldown assays, overnight cultures were back diluted 1:100 in 25 ml LB supplemented with 100 µg/mL ampicillin and grown at 37°C to an OD_600_ = 0.6. The cultures were induced using 0.3 mM IPTG and 200 µM C_6_HSL and incubated at 18°C with shaking at 130 rpm for 8 h. The cells were harvested at 4500 rpm for 10 min at 4°C and re-suspended in the ice-cold lysis buffer containing 50 mM Tris-HCl pH 8.0, 150 mM NaCl, 0.1% Triton X-100, and 7 mM imidazole. Cells were lysed using probe sonication (20% power, 15 sec on/off pulses for 3 min) and the cell lysates were spun at 14000 rpm for 30 min at 4°C to collect the soluble fraction. To check the solubility of WT or RhlR variants, whole cell lysates and soluble fractions were analyzed using SDS-PAGE. For affinity purification pulldowns, Ni-NTA magnetic beads (New England Biolabs) were incubated with soluble protein fractions. 30 µl pre-equilibrated magnetic beads with lysis buffer were added to 1 mL of each soluble fractions (input) and incubated at 4°C for 45 min with mixing. The protein bound beads were washed with 20-column volume of wash buffer (50 mM Tris-HCl pH 8.0, 150 mM NaCl, 0.1% Triton X-100, 30 mM imidazole). The bound protein complexes were collected using elution buffer (50 mM Tris-HCl pH 8.0, 150 mM NaCl, 0.1% Triton X-100, 300 mM imidazole) (output). To evaluate the interaction between PqsE and WT RhlR or RhlR variants, both input and output fractions were analyzed using SDS-PAGE. Gels were stained using Coomassie brilliant blue and imaged on a BioRad EZ-Doc gel imager.

For large scale purifications, 1 L of culture was used. RhlR and RhlR A44M were co-purified with 6x-His-PqsE using the same protocol described for the affinity purification pulldown method. The cells were harvested at 8000 rpm at 4°C, resuspended in 40 mL of lysis buffer and then lysed using probe sonication at 20% amplitude with a 15 sec on/off cycle for 15 min on ice. The cell lysate was spun at 14000 rpm at 4°C for 45 min to remove the cell debris. The supernatant was collected and then mixed with 1 mL of Ni-NTA beads (Qiagen) that were pre-equilibrated with lysis buffer. The slurry was incubated at 4°C for 45 min. The protein bound beads were washed with 15 mL of wash buffer and protein was eluted using the elution buffer. Ni-NTA fractions containing PqsE and RhlR were concentrated and further purified using a Superose-6 column (Cytiva) pre-equilibrated with 50 mM Tris HCl pH 8.0, 150 mM NaCl and 200 µM C_6_HSL. Protein purity was checked via SDS-PAGE. The gel was stained with Coomassie brilliant-blue and imaged on a Bio-Rad EZ-Doc gel imager.

### Pyocyanin production assay

For Figure 4A, overnight *P. aeruginosa* cultures were grown in 3 mL LB at 37°C. Pyocyanin was measured as previously described (34). For Figure 4B, overnight *P. aeruginosa* cultures were grown and back diluted 1:100 in 20 mL LB containing 2 μM C4HSL or a DMSO carrier control in strains lacking *rhlI*. Back dilutions were grown for 5 h at 37°C and pyocyanin was measured as previously described. For Figure 4C, 3 μL of overnight *P. aeruginosa* cultures were spotted on Congo red plates and grown for 5 days in the dark at RT. Pyocyanin was extracted from the entire plate using chloroform, then extracted into 0.1M HCl for quantification, and measured at 520_nm_ (56). For Figure 6D, pyocyanin production assays were performed by measuring pyocyanin from planktonic cultures supernatants. The PA14 WT and strains containing RhlR variants T58L and A44M were transformed with the pJN105 vector with or without the *pqsE* gene controlled by the pBAD promoter. The overnight cultures were washed and resuspended in LB media containing gentamycin (30 µg/ml) for plasmid selection and the secondary cultures were started at an A600_nm_ = 0.1. The cultures were incubated at 37°C with shaking at 200 rpm. At A600_nm_ = 0.6, the cultures were induced with a final concentration of 0.1% (v/v) arabinose. Cultures were incubated for 10 h and the culture supernatants were harvested by centrifugation. The cell-free supernatant was measured at A695_nm_.

### Swarming assay in *P. aeruginosa*

Swarming plates were made with a 5X M9 salt stock solution consisting of 2.7 g NH_4_Cl, 4.25 g Na_2_PO_4_, 7.5 g KH_2_PO_4_, and 1.25 g NaCl in 500 mL deionized water (57). 40 mL 5X M9 stock was added to 150 mL deionized water with 0.8 g agar for a final agar concentration of 0.4%. 200 µL of filter sterilized 1 M MgSO_4_, 200 µL filter sterilized 1 M CaCl_2_, and 2.7 g dextrose in 10 mL 0.5% casamino acids for a total concentration of 10%. 5 mL overnight cultures were grown and normalized to an OD_600_ = 1. 5 µL of normalized cell cultures were spotted on the center of the plate and left to dry. Plates are incubated face down at 37°C for 24 h and for another 12 h at room temperature. Images were taken using an Epson Perfection V850 Pro scanner at a resolution of 300 dpi and a display gamma of 2.2.

### Rhamnolipid production assay in *P. aeruginosa*

Clear 96-well plates (CellTreat) were dyed using 50 µL 0.1 mg/mL of Victoria Pure Blue BO in isopropanol (58). The isopropanol was evaporated under a vacuum without spinning for 1 h at 45°C. 300 µL of 0.5 M NaOH was added to the wells to fix the dye and incubated at room temperature for 10 min. The NaOH was aspirated, and the plates were dried under a vacuum without spinning for 1 h at 45°C. 5 mL overnight cultures were grown and back diluted into 1 mL cultures grown in 24-well plates (Greiner Bio-One) for 24 h. 1 mL of supernatant was collected after spinning the plate for 2 min at 15060 rpm. In triplicate, 250 µL supernatant and a rhamnolipid control were added to dyed wells and incubated with shaking at 750 rpm for 1 h at room temperature. 200 µL were transferred to a clean 96-well plate (CellTreat), and absorbance was read at 625_nm_ with a 5 s shake before reading (Molecular Devices SpectraMax M5 SoftMax Pro5).

### Congo red biofilm assay and *phz2* reporter expression in biofilms

Biofilm plates were made with 1% tryptone including 15.2 µM NH_4_SO_4_, 0.36 µM FeSO_4_, 1 mM KH_2_PO_4_ monobasic, 49 µM Congo red dye (w/v) (Sigma Aldrich), and 12.5 µM Coomassie blue dye (w/v) (Sigma Aldrich). Agar was added to a final concentration of 1%. 1 mM filter sterilized MgSO_4_, 0.2% (w/v) Casamino acids, and 0.5% (w/v) glycerol were added to cooled agar mixture after autoclaving prior to pouring. 5 mL overnight cultures expressing the *phz1/2* mScarlet transcriptional reporters were grown and 3 µL were spotted at the center of the plate and left to dry. Plates were incubated face down in an air-tight box to control for humidity at room temperature in the dark for 5 days. Brightfield and red fluorescent images were taken after 5 days.

### Image analysis

#### Swarming quantification

Traces on swarming plate images were segmented through manual thresholding (0–255), based on pixel value intensities of each image. For swarming distance analysis, from the user defined center point (x_0_, y_0_), the farthest distance traveled was calculated as the maximum value of distances of all boundary points to the center point (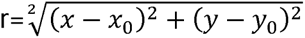).

#### Biofilm fluorescence signal analysis

Intensities of colony biofilms expressing the *phz1/2* mScarlet transcriptional reporters grown on Congo Red plates were corrected to WT UCBPP-PA14 non-fluorescence controls at each time point. For each corrected image, the sum of fluorescence intensity is calculated as ∑ *Pixel Value*. Preprocessing and calculation were carried out with functions in the scipy and cv2 packages, see https://github.com/biqingliang/Structural-Basis-for-C4-selection-by-PA14-RhlR for deposited scripts.

### Western blot analysis

#### E. coli

Strains were grown and induced as described previously (see ***E. coli* Light Assay**), with a culture size of 15 ml. Bacterial pellets were collected by centrifugation at 10,000 rpm for 10 minutes at 4°C and stored at -20°C until processed. Cell pellets were lysed in 1% SDS, 150 mM NaCl, and 50 mM Tris pH 8.0 for 5 min and subsequently sonicated for 15 s at 10% amplitude three times. Whole cell lysate was then subjected to Pierce BCA Protein Assay Kit (ThermoFisher) to determine protein concentration. Loading dye (250 mM Tris pH 6.8, 8% SDS, 40% glycerol, 8% 2-mercaptoethanol) was added to the lysates and boiled at 100°C for 5 min. Protein was loaded at a concentration of 20 µg/well into 4-20% Mini-PROTEAN TGX Precast Protein Gels (Bio-Rad) and run in a Mini-PROTEAN Tetra Vertical Electrophoresis Cell (Bio-Rad) at 200V for 45 minutes. Gels were transferred to activated (100% methanol, 1 min) PVDF (Millipore Sigma) using a Trans-Blot SD Semi-Dry Transfer Cell (Bio-Rad). Total protein images were acquired using a ChemiDoc MP (Bio-Rad). Membranes were blocked in 5% dry milk powder in TBST for 1 hour. Membranes were then incubated overnight in 5% milk in TBST with 1:1000 α-RhlR (10) and washed three times for 5 min in TBST prior to the addition of goat α-rabbit-HRP secondary (1:10000, Sigma). Chemiluminescent signal was detected using Pierce ECL Western Blotting Substrate (ThermoFisher) on a ChemiDoc MP. Band densitometry analysis was done in ImageJ (version 1.54m) (59) using total protein image for normalization.

#### P. aeruginosah

Strains were grown with or without AHL in a culture size of 20 mL until reaching an OD_600_ = 2.0. For the PqsE western blots, bacterial pellets were collected by centrifugation at 10,000 rpm for 10 minutes at 4°C. Cell pellets were lysed in 150 mM NaCl, 50 mM Tris pH 8.0-, and 20-mM imidazole and sonicated for 15 seconds at 25% amplitude three times. Whole cell lysate was then subjected to Pierce BCA Protein Assay Kit (ThermoFisher) to determine protein concentration. For the RhlR Westerns, loading dye (250 mM Tris pH 6.8, 8% SDS, 40% glycerol, 8% 2-mercaptoethanol) was added to the lysates and boiled at 100°C for 10 minutes. Protein was loaded at a concentration of 20 µg/well into 4-20% Mini-PROTEAN TGX Precast Protein Gels (Bio-Rad) and run in a Mini-PROTEAN Tetra Vertical Electrophoresis Cell (Bio-Rad) at 35 mA for 35 minutes. Gels were transferred to activated (100% methanol, 1 min) PVDF (Millipore Sigma) using a Trans-Blot SD Semi-Dry Transfer Cell (Bio-Rad). Membranes were blocked in 5% dry milk powder in TBST for 1 h. Membranes were then incubated overnight in 5% dry milk powder in TBST with 1:1000 α-PqsE. Membranes were washed three times for 15 min in TBST prior to the addition of goat α-rabbit-HRP secondary (1:10000, Sigma) in 5% dry milk powder in TBST. Chemiluminescent signal was detected using Pierce ECL Western Blotting Substrate (ThermoFisher) on a ChemiDoc MP.

### *rhlA* promoter-fusion activity assay in *P. aeruginosa*

The *rhlA* promoter mScarlet fusion construct was cloned in pUCP18 using restriction enzyme cloning (see Table S2). The construct was electroporated in PA14 WT, Δ*rhlI* and Δ*rhlR* strains. Primary overnight cultures were grown at 37°C and secondary cultures were back diluted 1:100 in 5 mL LB media supplemented with 400 µg/mL carbenicillin. The cultures were incubated for 24 h at 37 °C with shaking at 200 rpm. 1 mL of culture was pelleted at 3000 rpm for 5 min. The supernatant was discarded, and the pellet was resuspended in 1 mL of 1x PBS. Fluorescence was recorded using a Glomax (Promega) plate reader with excitation at 520 nm and emission at 540-580 nm in a clear bottom black opaque 96-well plate (Corning).

### RNA extraction and RNA-seq data analysis

#### RNA extraction

5 mL overnight cultures were back diluted 1:100 in 20 mL of fresh LB media and incubated for 5 h at 37°C with shaking. Cells were harvested by centrifugation and pellets frozen until RNA extraction. RNA was extracted using the Invitrogen™ PureLink™ RNA Mini Kit. Pellets were resuspended in 100 µL lysozyme solution containing 10 mM Tris-HCl pH 8.0, 0.1 mM EDTA, and 1 mg/mL lysozyme. 5 µL of 1% SDS was added, the solution was mixed with a vortex and incubated at room temperature for 5 min. 350 µL lysis buffer (Invitrogen) was added with 10 µL 2-mercaptoethanol for each 1 mL Lysis Buffer and vortexed. Cells were lysed by bead beating (BioSpec 0.1 MM Zirconia/Silica Beads) for two rounds of 50 s. Lysed cells were transferred to fresh microcentrifuge tubes with 250 µL of 100% ethanol and mixed with a vortex. Samples were spun in spin cartridges (Invitrogen) at 12,000 x g for 15 s at room temperature, and the flow-through was discarded. 700 µL Wash Buffer I (Invitrogen) was added and spun at 12,000 x g for 15 s at room temperature, and the flow-through was discarded. 500 µL Wash Buffer II (Invitrogen) with ethanol was added and centrifuged for 12,000 x g for 15 s at room temperature; the flow-through was discarded, and RNA samples were washed again. Cells were spun at 12,000 x g for 1 min at room temperature to dry the membrane with the bound RNA. 50 µL RNAse free water was added to the center of the spin cartridge, incubated at room temperature for 1 min, and centrifuged for 2 min at ≥ 12,000 x g at room temperature. Extracted RNA was then frozen at -80°C until further use.

#### Data analysis

Bulk RNA was sent to SeqCenter (Pittsburgh, PA) where it was processed for paired-end Illumina sequencing via rRNA depletion using Ribo-Zero Plus (Illumina). Fastq files generated by SeqCenter were processed using Rockhopper with default settings (60–62). Briefly, fastq files were aligned to the genome of *Pseudomonas aeruginosa* UCBPP-PA14. The reads were subsequently normalized using upper quartile normalization and transcript abundance was determined by reads per kilobase per million mapped reads (RPKM). Differential gene expression analysis was conducted using a negative binomial distribution model, outputting a p-value for each transcript. The p-values were then corrected for false-discovery using the Benjamini-Hochberg procedure, resulting in the q-values reported here (Table S2). SignalMap v2.0 (NimbleGen Systems) was used to visualize read counts for *phzABCDEFG1/2*, *phzH*, *phzM*, and *phzS* (Figure 5). Aligned sequences generated in Rockhopper were converted to GFF files and genes of interest were isolated into unique GFF files (37).

### Liquid chromatography mass spectrometry of QS metabolites

#### Molecule extraction

5 mL overnight cultures were back diluted 1:100 in 20 mL fresh LB media containing 2 μM C4HSL or a DMSO carrier control in strains lacking *rhlI*. Back dilutions were grown for 5 h at 37°C, supernatants were harvested by centrifugation, and frozen until use. Sample supernatants were thawed and 200 μL were added to a new 1.5 mL centrifuge tubes. Samples were extracted with 800 μL methanol, vortexed for 15 s, and centrifuged for 2 min at 10,000 x g. Samples were mixed 1:1 with 0.1% formic acid in an HPLC vial and vortexed again.

#### Analytical measurements and analyses

An ultra-high-performance liquid chromatography (UHPLC) system coupled with a high-resolution mass spectrometer (HRMS) was employed for analyte quantification. The analytical platform consisted of a Vanquish UHPLC system interfaced with a Thermo LTQ Orbitrap mass spectrometer (Thermo Fisher Scientific), operating in positive electrospray ionization (ESI) mode. Chromatographic separation was performed on a Hypersil GOLD C18 column (50 × 2.1 mm, 1.9 μm particle size; Thermo Fisher Scientific) using mobile phase A (0.1% v/v formic acid with 5 mM ammonium formate in water) and mobile phase B (0.1% v/v formic acid with 5 mM ammonium formate in methanol). The flow rate was set at 0.45 mL/min, and the column temperature was maintained at 45 °C. The gradient elution program started at 5.0% B (held for 1 min), increased linearly to 100% B over 3.5 min, held at 100% B for 1 min, then returned to 5.0% B over 0.1 min and held for an additional 1.4 min to re-equilibrate the column. The injection volume was 10 μL. To enable accurate quantification, a series of external controls at varying concentrations were run at the same batch with the unknown samples to generate a calibration curve. Unknown sample signals were normalized to this calibration curve to correct for batch-specific variability and instrument response drift. Quantification was performed using a 0.01 Da accurate mass window for peak integration. All data were acquired and processed using Xcalibur software (version 2.1, Thermo Fisher Scientific).

### Chromatin Immunoprecipitation (ChIP) Sequencing

*P. aeruginosa* strains were grown overnight, back diluted 1:100 in fresh LB media and grown to an OD_600_ = 2.0. Crosslinking, lysis, and immunoprecipitation were performed as previously described (37).

### ChIP-qPCR

2 mL overnight cultures were back diluted 1:100 in 20 mL fresh LB media and incubated at 37°C with shaking until they reached an OD_600_ = 2. Cells were crosslinked by adding 500 µL formaldehyde for a final concentration of 1%, were swirled to mix, and left for 20 min. Crosslinked cultures were transferred to 50 mL conical tubes and quenched with 10 mL 2.5 M glycine for a final concentration of 0.5 M. Cells were pelleted at 4000 rpm for 5 min, and the supernatant was discarded. Pellets were resuspended in 20 mL 1X TBS and spun at 4,000 rpm for 5 min. The supernatant was discarded, the pellet resuspended in 1 mL 1X TBS, and cells were transferred to a microcentrifuge tube. Cells were spun at 14,000 rpm for 5 min, the supernatant was discarded, and the cell pellets were frozen until further use. Thawed cells were resuspended in 1 mL 150 mM FA Lysis Buffer (1 M HEPES, pH 7.5, 2 M sodium chloride, 0.5 M EDTA, 100% Triton-X 100, 5% sodium deoxycholate, 10% SDS, water) containing 4 mg/mL lysozyme (Sigma) and incubated at 37°C for 30 min to lyse cells. Lysed cells were then sonicated at 0-10°C for 30 min with 30 s on/off pulses at 75% output with a BioRuptor Sonicator (Diagenode). Cell lysates were centrifuged twice for 10 min at 14,000 rpm. Between each centrifugation, supernatant was transferred to new microcentrifuge tubes. 1 mL cell lysate supernatants were diluted in 1 mL 150 mM FA Lysis Buffer for a final volume of 2 mL. 100 µL cross-linked, sonicated lysates were diluted in 800 µL 150 mM FA Lysis Buffer, and 20 µL were saved as input samples. 30µL of a 50% slurry of protein A beads in 1X TBS (Sigma) and 10 µL RhlR antibody were added to lysates and incubated at room temperature for 90 min on a rotisserie rotator. The immunoprecipitation mix was centrifuged for 1 min at 4,000 rpm, the supernatant was removed by aspiration, and the pellet was resuspended in 750 µL 150 mM FA Lysis Buffer. This mix was transferred to a Spin-X column (Corning), incubated for 3 minutes at room temperature on a rotisserie rotator, spun again for 1 minute at 4,000 rpm, and the supernatant was removed by aspiration. This was repeated with 700 µL 150 mM FA Lysis Buffer, 500 mM FA Lysis Buffer (1 M HEPES, pH 7.5, 2 M sodium chloride, 0.5 M EDTA, 100% Triton-X 100, 5% sodium deoxycholate, 10% SDS, water), ChIP Wash Buffer (1 M Tris-HCl, pH 8.0, 1 M lithium chloride, 0.5 M EDTA, NP-40, 5% sodium deoxycholate), and TE (10 mM Tris-HCl pH 8.0, 1 mM EDTA). After the last wash, protein A beads were resuspended in 100 µL ChIP Elution Buffer (1 M Tris-HCl pH 7.5, 0.5 M EDTA, 10% SDS), the Spin-X column was placed in a fresh dolphin-nosed tube (Sorenson) and incubated for 10 min at 65°C with occasional gentle mixing. Immunoprecipitation mixes were centrifuged for 1 min at 4,000 rpm, and the Spin-X filters were discarded. Cells and input samples were de-crosslinked by incubating for 10 minutes at 100°C in microcentrifuge tubes. DNA was cleaned up with a PCR purification kit (Qiagen). Input samples were diluted 1:100 for 10 µL qPCR reactions. qPCR reactions were set up with 5 µL SYBR Select Master Mix (ThermoFisher), 2 µL cleaned DNA, and 3 µL combined forward and reverse primers. qPCR reactions were run with a 50°C holding stage, a 95°C holding stage, 40 cycles of 95°C to 60°C, and a continuous melt curve stage from 95°C to 60°C (Applied Biosystems 7500 Fast Real-Time PCR System).

### Electrophoretic mobility shift assays (EMSA)

EMSAs were performed with promoter regions of *rhlA* and *phzA1* gene as previously described with minor modifications. To obtain the respective DNA fragment, 320 bp upstream of *rhlA* gene and 723 bp upstream of *phzA1* gene was PCR amplified from genomic DNA of *P. aeruginosa* PA14 strain. Concentrations of PqsE, RhlR:C_6_HSL bound to PqsE, RhlR A44M:C_6_HSL bound to PqsE normalized per reaction. The EMSA reaction contained of 1 µl of 10 ng/µL of the promoter DNA fragment, 4 µl of protein dilution (0.9375 µM, 1.875 µM, 3.75 µM, 7.5 µM, 15 µM) and 15 µL of EMSA buffer (20 mM Tris-HCl pH 7.5, 30 mM NaCl, 0.5 mM EDTA, 0.2 mg/mL BSA, 10% glycerol, 1 mM DTT, 10 mM MgCl_2_). Reactions were mixed by vortex and incubated at 30°C for 15 min. 8 µl of the completed reaction was mixed with Novex Hi-Density TBE 5X Sample Buffer (Invitrogen) and loaded on an 8% polyacrylamide gel. Electrophoresis was conducted in a 1X TBE buffer at 100 V for 60 min at 4°C. The gels were washed using a 0.5X TB buffer. The gels were stained using 1X gel red (GoldBio) in 50 mL of 0.5X TB buffer with shaking followed by three washes with 50 mL 0.5X TB buffer. Imaging was performed on Bio-Rad EZ-Doc gel imager using a UV sample tray capable of capturing emission spectrum of 550-700 nm (Bio-Rad) compatible with GelRed DNA staining dye.

## DATA AVAILABILITY

Sequencing data have been deposited at the NCBI Sequence Read Archive under the submission number PRJNA1338225. Code related to the imaging analyses performed in the manuscript can be found at https://github.com/biqingliang/Structural-Basis-for-C4-selection-by-PA14-RhlR.

## SUPPORTING INFORMATION

**Table S1.** Read counts for all mRNA from RNA-seq experiments for all strains.

**Table S2.** Strains and plasmids used in this study.

**Table S3.** Oligonucleotides used in this study.

**Figure S1. Expression of RhlR variants in the *E. coli* reporter assay.** WT RhlR and RhlR variants expressed from the pBAD-A vector in *E. coli* as measured by western blot analysis using a polyclonal antibody for RhlR. An empty vector (EV) control was used to assess antibody specificity.

**Figure S2. Structural analysis of the RhlR LBP.** Zoom in view of the RhlR (pink) LBP. C_6_HSL (salmon), which was computationally docked into the WT RhlR LBP (PDB: 8DQ0) (top) and mBTL (green) from experimental results (PDB: 8DQ0) are shown. Residues A44, G46, T58, Y64, W68, L69, Y72, D81, I84, W96, W108, and V133 are shown as sticks. Residues are colored based on variant responses to C_4_HSL: orange = substitutions are recalcitrant to C_4_HSL; magenta = substitutions are hypersensitive to C_4_HSL; gray = substitutions are hyposensitive to C_4_HSL; dark green = phenylalanine substitutions were previously described and lead to a “constitutive” state for RhlR.

**Figure S3. RhlR G46 variants are less responsive to C_4_HSL than WT RhlR.** RhlR-controlled bioluminescence was measured in *E. coli*. Arabinose-inducible RhlR was expressed from one plasmid and a p*rhlA*-*luxCDABE* reporter construct was carried on a second plasmid to monitor transcriptional activity. 0.1% arabinose was used to induce RhlR. RhlR-dependent bioluminescence was measured for WT RhlR (black) and RhlR variants (G46M = blue, G46W = purple) in response to increasing concentrations (µM) of C_4_HSL with or without PqsE.

**Figure S4. RhlR L69/I84/W108 variants are less responsive to C_4_HSL than WT RhlR.** RhlR-controlled bioluminescence was measured in *E. coli*. Arabinose-inducible RhlR was expressed from one plasmid and a p*rhlA*-*luxCDABE* reporter construct was carried on a second plasmid to monitor transcriptional activity. 0.1% arabinose was used to induce RhlR. RhlR-dependent bioluminescence was measured for WT RhlR (black) and RhlR variants**; top)** (L69D = blue, L69K = purple, L69M = orange), **middle)** (I84M = blue, I84W = purple), and **bottom)** (W108F = blue, W108Y = purple) in response to increasing concentrations (µM) of C_4_HSL with or without PqsE.

**Figure S5. RhlR Y72/D81/W96 variants are non-responsive to C_4_HSL than WT RhlR.** RhlR-controlled bioluminescence was measured in *E. coli*. Arabinose-inducible RhlR was expressed from one plasmid and a p*rhlA*-*luxCDABE* reporter construct was carried on a second plasmid to monitor transcriptional activity. 0.1% arabinose was used to induce RhlR. RhlR-dependent bioluminescence was measured for WT RhlR (black) and RhlR variants; **top)** (Y72A = blue, Y72T = purple), **middle)** (D81K = blue, D81N = purple), and **bottom)** (W96A = blue) in response to increasing concentrations (µM) of C_4_HSL with or without PqsE.

**Figure S6. Purification of WT RhlR:C_6_HSL-PqsE and RhlR A44M-PqsE.** Representative SDS-PAGE gel of whole-cell lysate (W), supernatant (S), and elution (E) for Ni-NTA purification of WT RhlR:C_6_HSL-PqsE as well as fractions 11-15 (Figure 3C) from Superose-6 size-exclusion chromatography for RhlR:C6HSL-PqsE and RhlR A44M-PqsE.

**Figure S7. RhlR T58V/L and A44M variants are expressed and differentially regulate *rhlA* and swarming. A)** Expression of WT RhlR and RhlR variants from its native locus in *P. aeruginosa* as measured by western blot analysis using a polyclonal antibody for RhlR. **B)** Colony forming units of cells collected from colony biofilms for pyocyanin extraction shown in Figure 4C, 4D. **C)** Branch length measurements of strains expressing WT RhlR and RhlR variants compared to Δ*rhlI* grown on swarming media. Branch length was measured using ImageJ by establishing the center of the initial inoculant and measuring in a direct line to the tip of the branch. **D)** Expression of plasmid-borne a p*rhlA*-mSacrlet reporter in WT RhlR and RhlR variants compared to Δ*rhlI*. Statistical analyses for all assays were performed using an ordinary one-way ANOVA with a Tukey’s multiple comparisons test; comparisons that were deemed not significant by these analyses are not shown. **E)** WT *P. aeruginosa* strains expressing WT RhlR (black), RhlR T58L (purple) and RhlR A44M (green) with the *phz1* (circle) and *phz2* (square) promoters fused to mScarlet integrated at the *attB* locus.

**Figure S8. Differential phenazine expression in biofilms expressing RhlR variants. A)** Representative brightfield and fluorescent images of Congo red colony biofilm plates for strains of *P. aeruginosa* expressing WT RhlR, RhlR T58L, and RhlR A44M with the *phz2* promoter fused to *mScarlet* integrated at the *attB* locus. Red signal indicates the expression of *phz2*-*mScarlet*. Images were merged using ImageJ. **B)** Quantification of fluorescent signal from three independent replicates from **A)** using a non-fluorescent strain of PA14 as a control. Statistical analyses for all assays were performed using an ordinary one-way ANOVA with a Tukey’s multiple comparisons test. Comparisons that were deemed not significant by these analyses are not shown.

**Figure S9. Standards and controls for LC-MS experiments.** Calibration standards for C_4_HSL, pyocyanin, phenazine-1-carboxylic acid (PCA), and phenazine-1-carboxamide (PCN) performed at 10, 31.6, 100, 316, 1000, 3160 nM. The area of the peak was measured using Xcalibur and plotted as a function of molecule concentration. A simple linear aggression analysis was calculated for each curve to determine the slope, which was used to calculate the values determined in Figure 5C.

**Figure S10. RhlR variant promoter occupancy.** Occupancy plot profiles showing the level of mapped ChIP-seq reads +/- 500 bp of the called peak centers for the designated strains at the **A)** *phzA1* and *phzM*, **B)** *rhlA* and **C)** *lecB* promoters. **D)** ChIP-qPCR analysis of WT RhlR and RhlR variant promoter binding to the regions upstream of *lecB* and *rhlA* to measure the relative levels of RhlR binding to DNA sites in a Δ*rhlI* strain containing 2 μM C_4_HSL. All data were normalized to a non-specific DNA control site.

## AUTHOR CONTRIBUTIONS

A.N.P., A.C., A.G.M, J.T.W. and J.E.P. conceived the study and designed experiments. A.N.P. performed experiments shown in Figures 2, 4, 5, 6, S3, S4, S5. V.R.B. performed experiments shown in Figures 3, 6, and S6. M.L.S. performed experiments shown in Figures 5, S1, and S6. A.G.M. performed the biofilm experiments shown in Figures 4, 5 and S8. A.M.S. provided technical assistance with the ChIP-seq experiment in Figure 6. C.P.M. conducted bioinformatic analyses displayed in Figure 1 and performed experiments shown in Figure 3. A.C. and A.F.K provided technical assistance for experiments shown in Figure 2 and Figure 4, respectively. B.L. performed the image analyses in Figure S7 and S8. J.E.P. and A.C. performed structural analyses of the RhlR LBP in Figures 1 and S2. X.E. provided technical expertise for experiments shown in Figure 5C and S7. A.N.P., V.R.B., M.L.S., C.P.M., A.G.M, X.K., J.T.W, and J.E.P. wrote the manuscript. A.N.P., V.R.B., M.L.S., C.P.M., A.C., A.F.K., J.T.W., and J.E.P. edited the manuscript. J.T.W and J.E.P. supervised the project.

## ACKNOWLEDGMENTS

The authors thank all members of the Paczkowski lab as well as the research laboratories in the Division of Genetics at the Wadsworth Center for helpful discussions on the research. The authors thank the Owen and Bartlett labs for sharing key resources. The authors thank the Rensselaer Polytechnic Institute Center for Biotechnology and Interdisciplinary Studies for technical support during the mass spectrometry experiments. This work was made possible with the help of the dedicated staff scientists at the Advanced Genomics Technologies Center and Media & Tissue Core facilities at the Wadsworth Center.

## FUNDING AND ADDITIONAL INFORMATION

This work was supported by National Institutes of Health training grant T32GM132066 to C.P.M., NIH grant R35GM144328 to J.T.W., NIH grant R01GM14436101, New York Community Trust Foundation grant P19-000454, Cystic Fibrosis Foundation grant PACZKO21G0, and American Lung Association Innovation Award INALA2023 to J.E.P. A.F.K. is supported by Cooperative Agreement Number NU60OE000104 (CFDA #93.322), funded by the Centers for Disease Control and Prevention (CDC) of the US Department of Health and Human Services (HHS).

## CONFLICT OF INTEREST

The authors declare no competing interests.

## ABBREVIATIONS AND NOMENCLATURE

AHL: acyl homoserine lactone
QS: quorum sensing
LBP: ligand binding pocket
PCA: phenazine-1-carboxylic acid
PCN: phenazine-1-carboxamide

## UNPUBLISHED OBSERVATIONS AND PERSONAL COMMUNICATIONS

N/A

## Notes

### Competing Interest Statement

The authors have declared no competing interest.

