## Supplementary figures and images for "RhlR quorum-sensing receptor ligand sensitivity regulates the differential expression of phenazine genes in *Pseudomonas aeruginosa*"

### Supplemental Figures

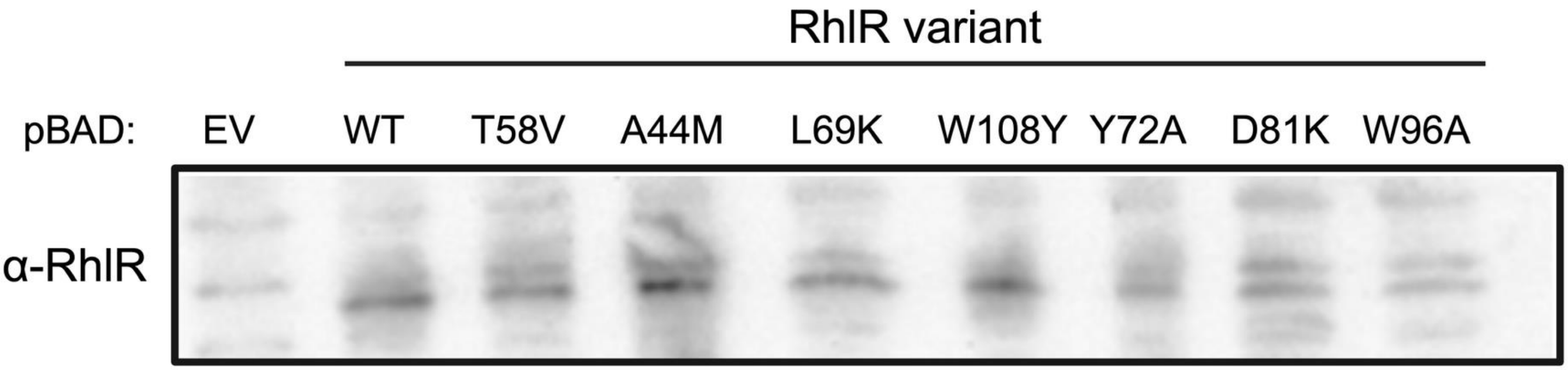

**Figure S2**

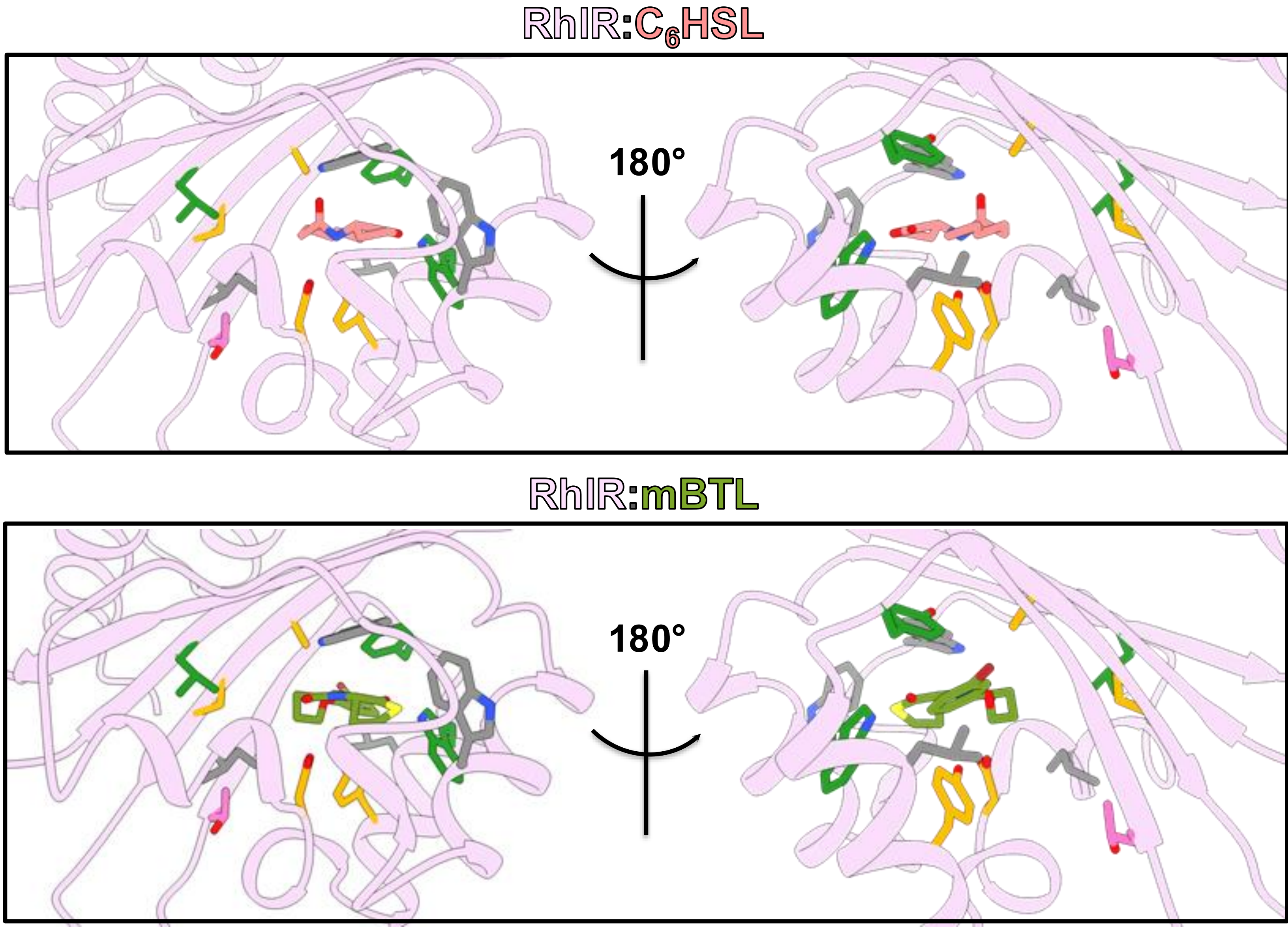

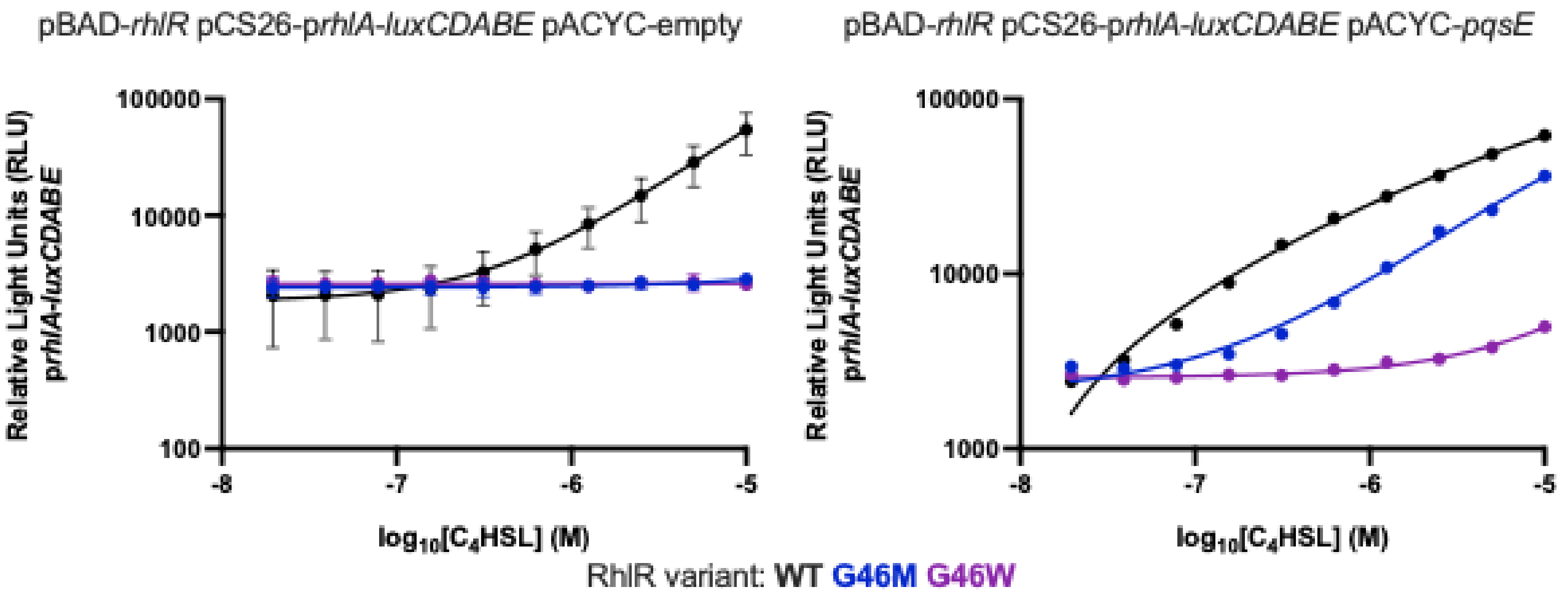

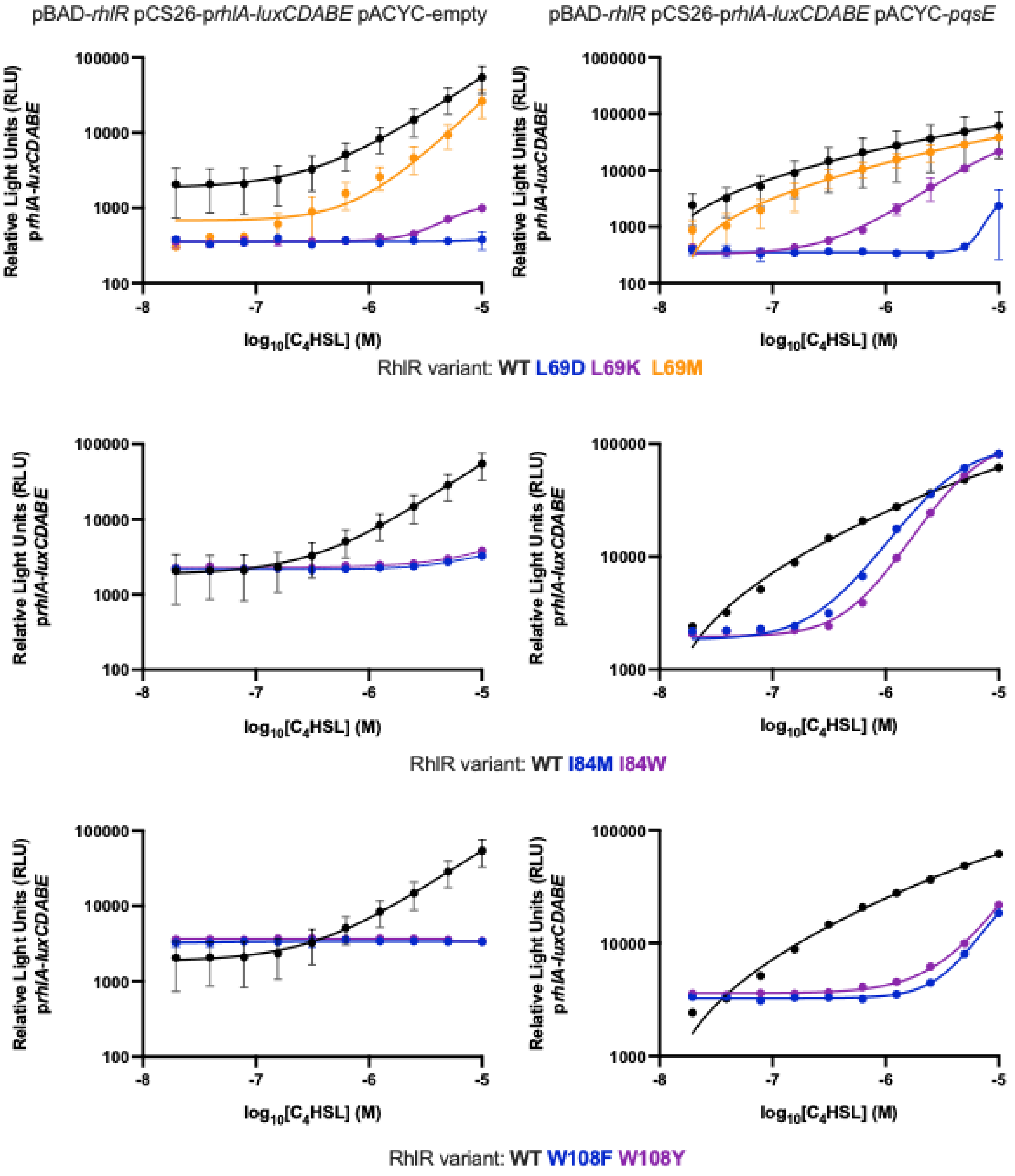

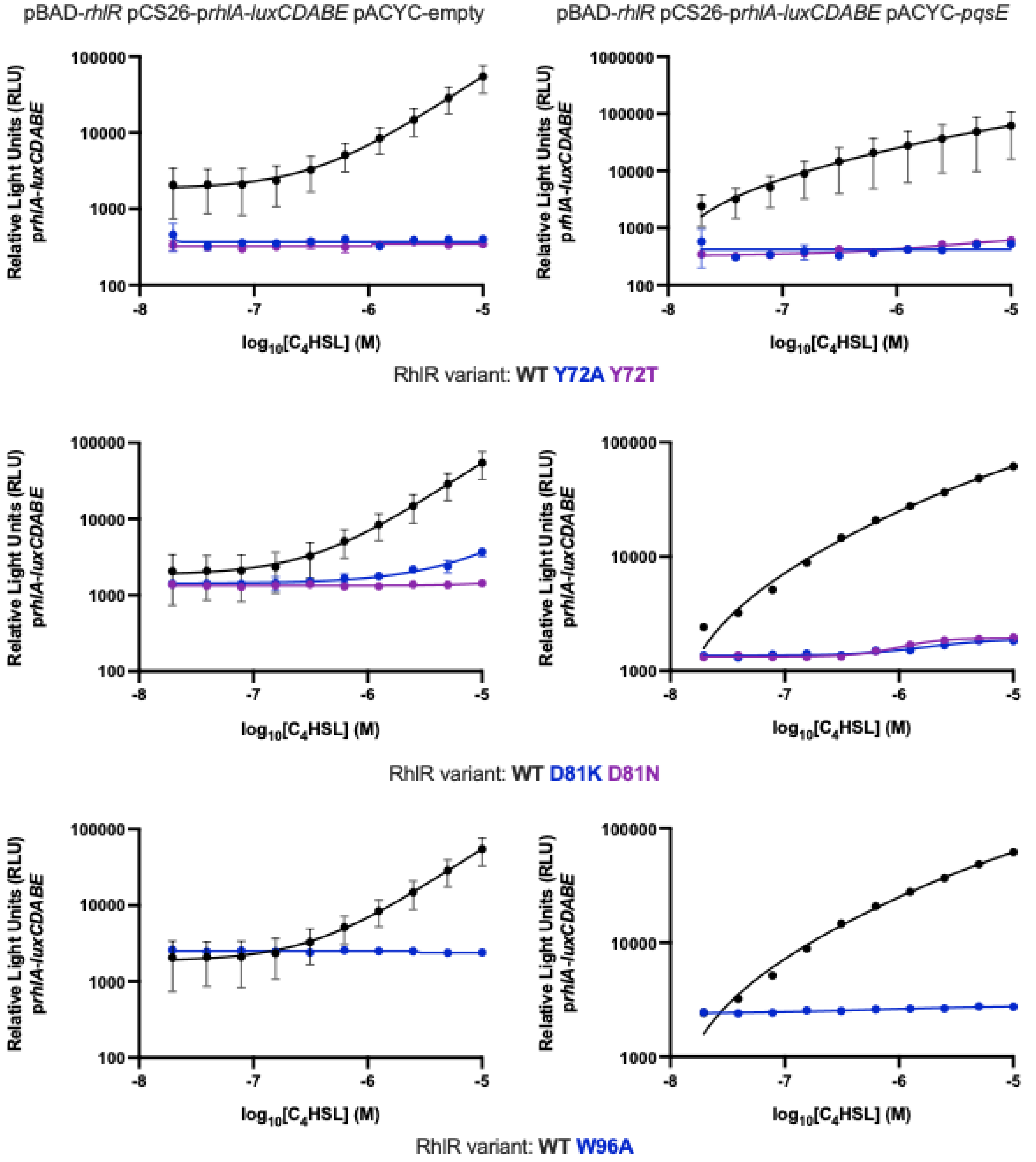

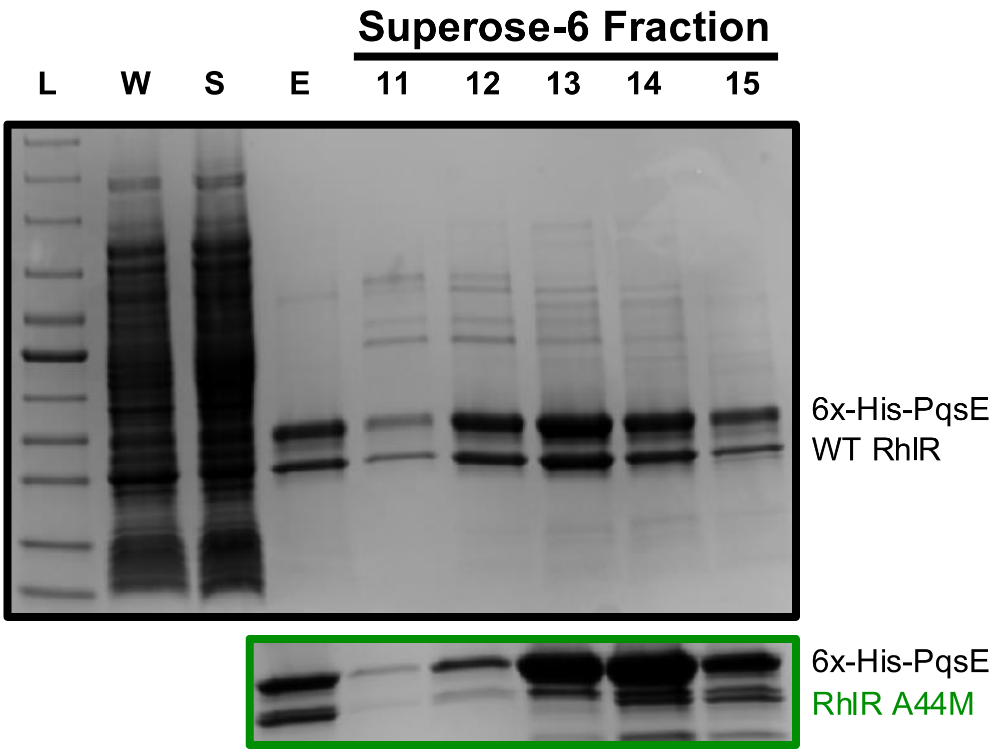

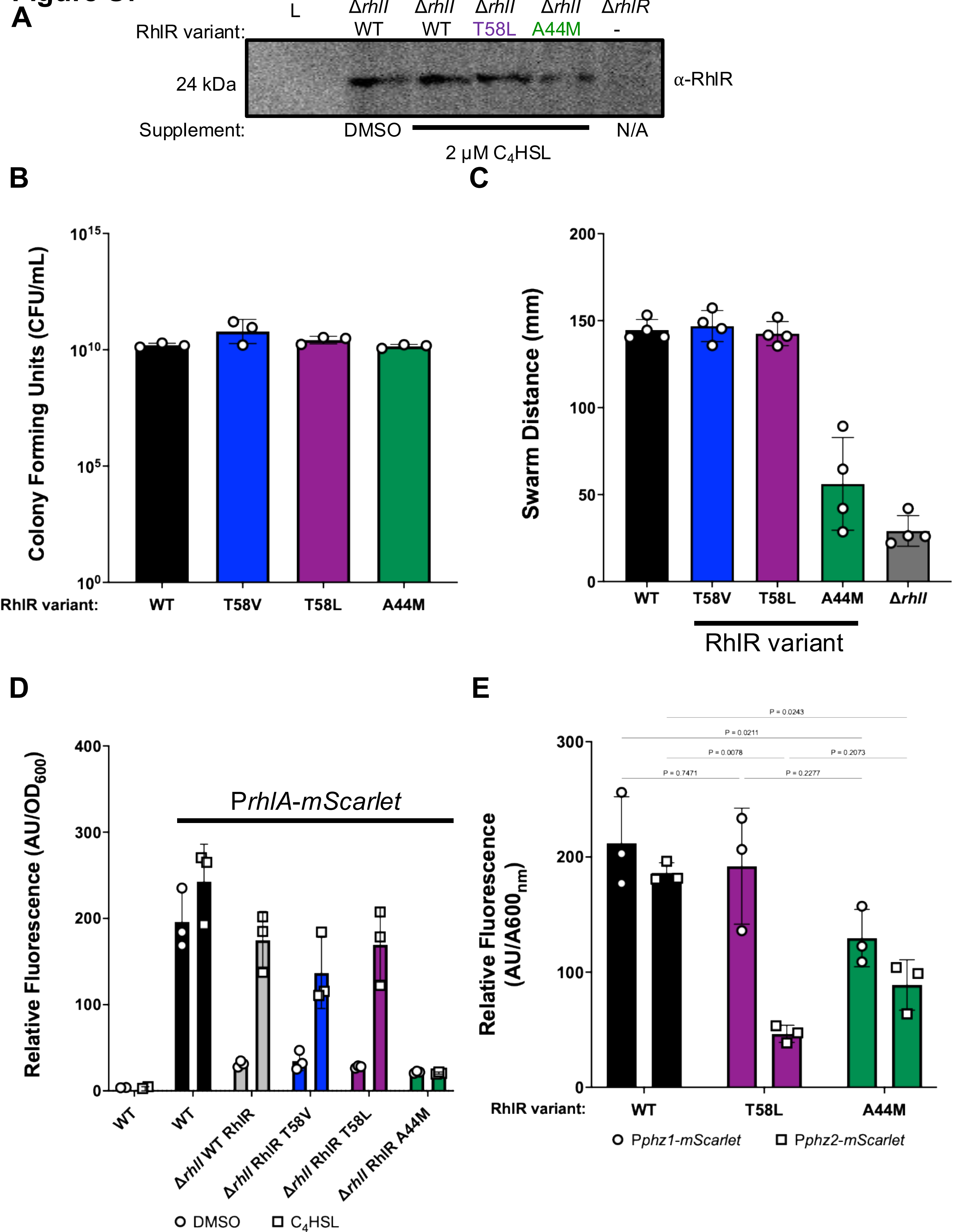

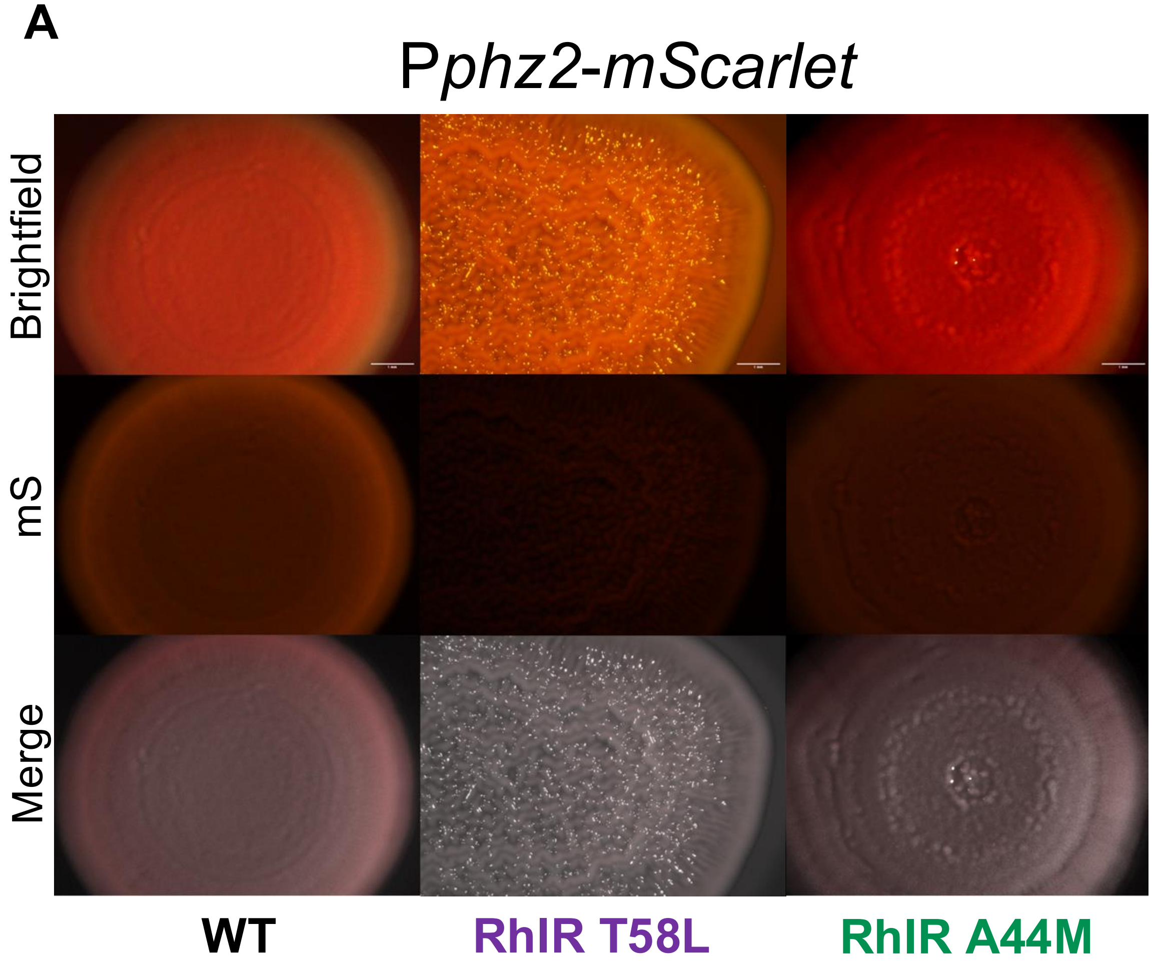

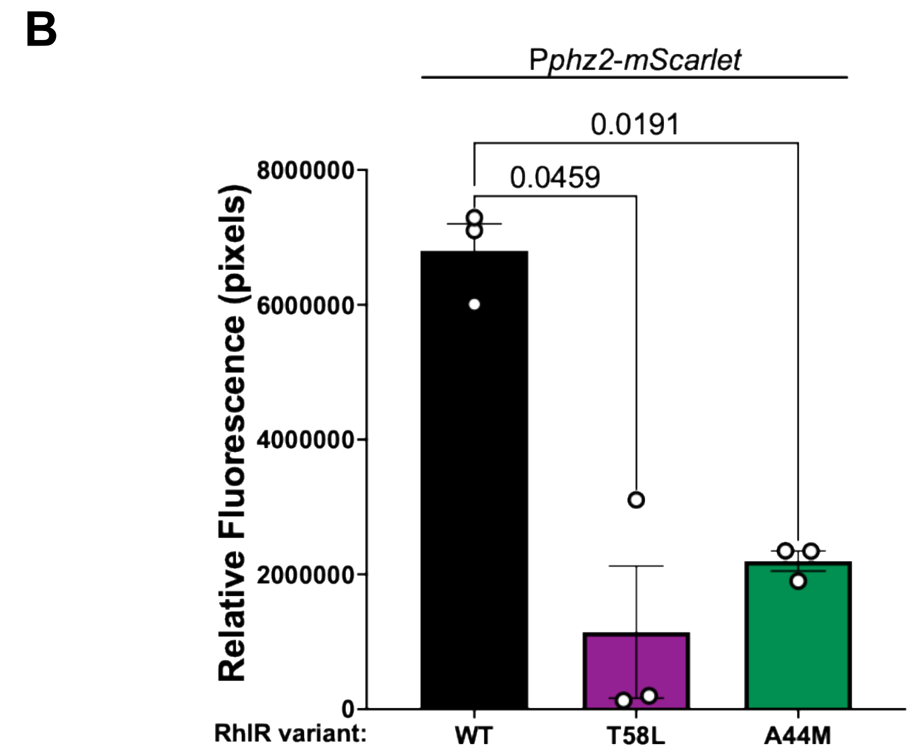

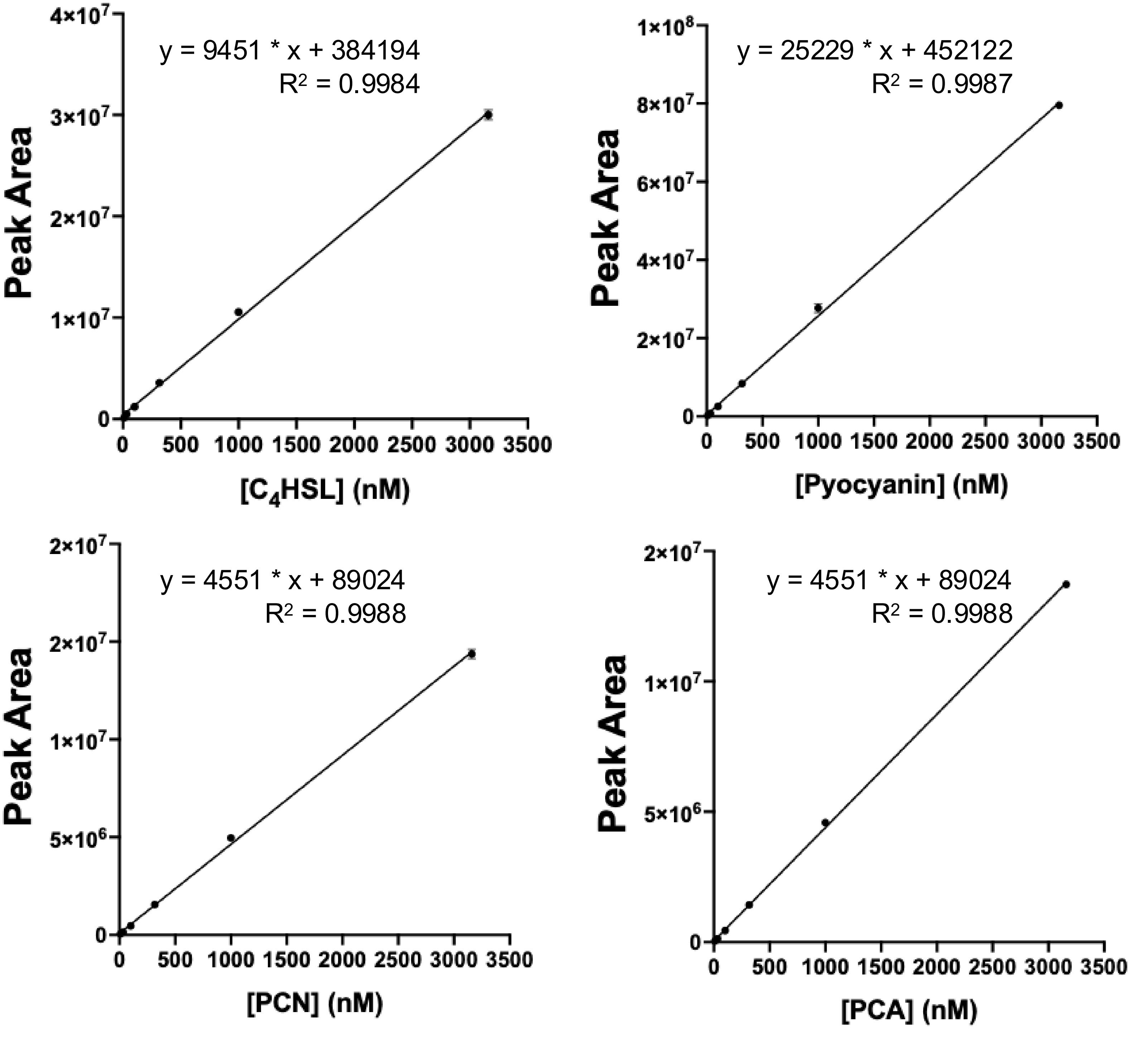

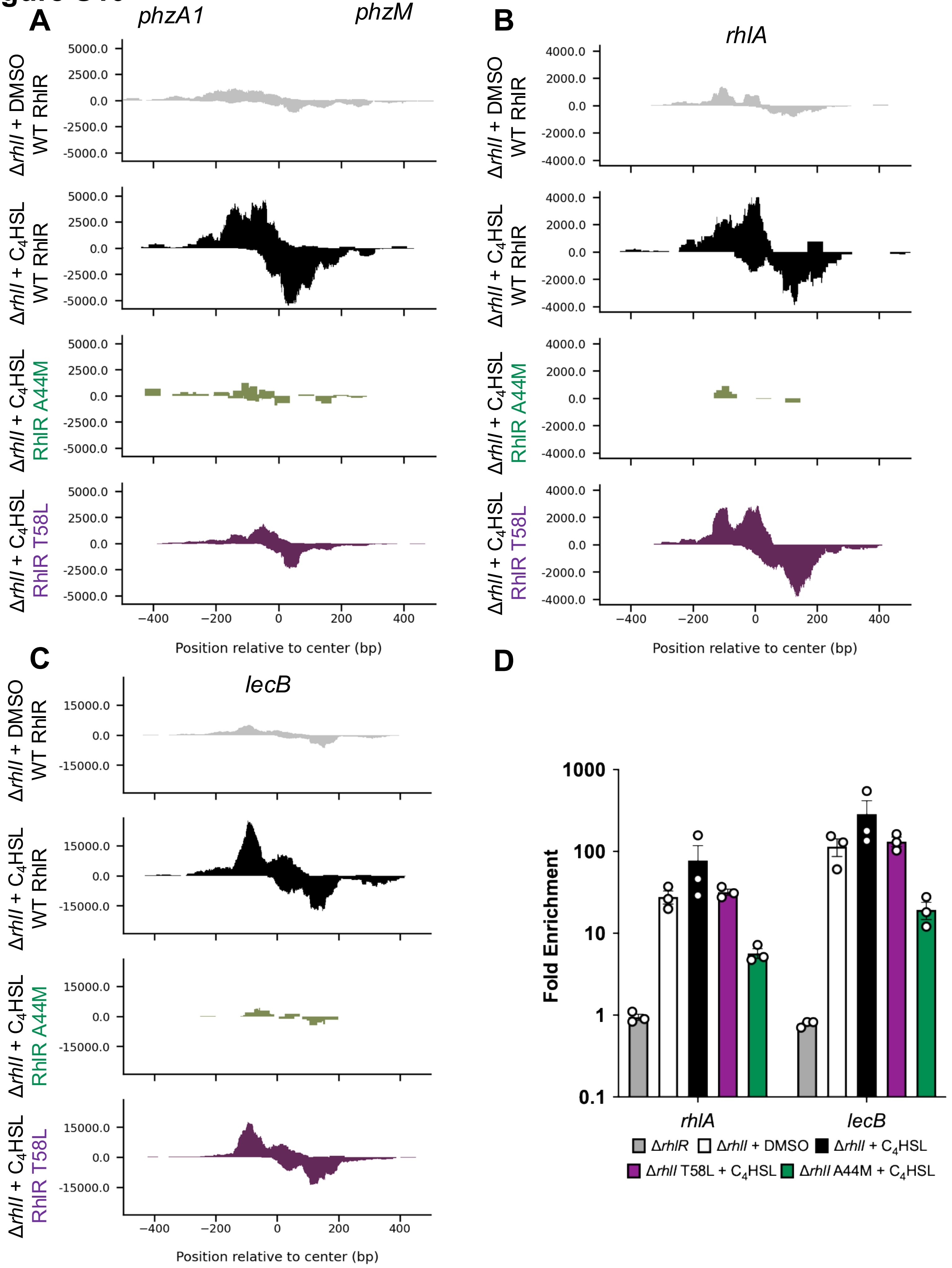
