## Supplementary material for "RhlR quorum-sensing receptor ligand sensitivity regulates the differential expression of phenazine genes in *Pseudomonas aeruginosa*": Table S2

**Table S2.** Strains and plasmids used in this study.

| **Strain #** | **Strain and Plasmid** | **Resistance** | **Source** |
| --- | --- | --- | --- |
| JPS0154 | PA14 Δ*rhlI* |  | Mukherjee *et al.* (2017) |
| JPS0222 | UCBPP-PA14 WT |  | Gift from Dr. George O'Toole |
| JPS0225 | *Escherichia coli* DH5α pETDuet | Amp | Invitrogen |
| JPS0226 | *E. coli* DH5α pBADA | Amp | Invitrogen |
| JPS0227 | *E. coli* DH5α pEXG2 | Gent | Gift from Dr. Joseph Mougous |
| JPS0476 | *E. coli* TOP10 pBAD-*rhlR*/pCS26-*rhlA*-*luxCDABE*/pACYC-*pqsE* | Amp/Kan/Tet | Taylor *et al.* (2021) |
| JPS0477 | *E. coli* TOP10 pBAD-*rhlR*/pCS26-*rhlA*-*luxCDABE*/pACYC-empty | Amp/Kan/Tet | Taylor *et al.* (2021) |
| JPS0961 | *E. coli* TOP10 pBAD-*rhlR* (Y72A)/pCS26-*rhlA*-*luxCDABE*/pACYC-*pqsE* | Amp/Kan/Tet | This study |
| JPS0962 | *E. coli* TOP10 pBAD-*rhlR* (Y72A)/pCS26-*rhlA*-*luxCDABE*/pACYC-empty | Amp/Kan/Tet | This study |
| JPS0963 | *E. coli* TOP10 pBAD-*rhlR* (T58V)/pCS26-*rhlA*-*luxCDABE*/pACYC-*pqsE* | Amp/Kan/Tet | This study |
| JPS0964 | *E. coli* TOP10 pBAD-*rhlR* (T58V)/pCS26-*rhlA*-*luxCDABE*/pACYC-empty | Amp/Kan/Tet | This study |
| JPS1179 | WT PA14 pUCP18-prhlA-mScarlet | Carb | This study |
| JPS1181 | PA14 Δ*rhlI* pUCP18-[rhlA-mScarlet | Carb | This study |
| JPS1283 | *E. coli* TOP10 pBADA-*rhlR* (Y72T)/pCS26-*rhlA*-*luxCDABE*/pACYC-*pqsE* | Amp/Kan/Tet | This study |
| JPS1284 | *E. coli* TOP10 pBADA-*rhlR* (L69K)/pCS26-*rhlA*-*luxCDABE*/pACYC-*pqsE* | Amp/Kan/Tet | This study |
| JPS1285 | *E. coli* TOP10 pBADA-*rhlR* (L69M)/pCS26-*rhlA*-*luxCDABE*/pACYC-*pqsE* | Amp/Kan/Tet | This study |
| JPS1286 | *E. coli* TOP10 pBADA-*rhlR* (L69D)/pCS26-*rhlA*-*luxCDABE*/pACYC-pqsE | Amp/Kan/Tet | This study |
| JPS1287 | *E. coli* TOP10 pBADA-*rhlR* (Y72T)/pCS26-*rhlA*-*luxCDABE/*pACYC-empty | Amp/Kan/Tet | This study |
| JPS1288 | *E. coli* TOP10 pBADA-*rhlR* (L69K)/pCS26-*rhlA*-*luxCDABE*/pACYC-empty | Amp/Kan/Tet | This study |
| JPS1289 | *E. coli* TOP10 pBADA-*rhlR* (L69M)/pCS26-*rhlA*-*luxCDABE*/pACYC-empty | Amp/Kan/Tet | This study |
| JPS1290 | *E. coli* TOP10 pBADA-*rhlR* (L69D)/pCS26-*rhlA*-*luxCDABE*/pACYC-empty | Amp/Kan/Tet | This study |
| JPS1464 | *E. coli* TOP10 pBADA-*rhlR* (I84M)/pCS26-*rhlA*-*luxCDABE*/pACYC-*pqsE* | Amp/Kan/Tet | This study |
| JPS1465 | *E. coli* TOP10 pBADA-*rhlR* (I84M)/pCS26-*rhlA*-*luxCDABE*/pACYC-empty | Amp/Kan/Tet | This study |
| JPS1466 | *E. coli* TOP10 pBADA-*rhlR* (I84W)/pCS26-*rhlA*-*luxCDABE*/pACYC-*pqsE* | Amp/Kan/Tet | This study |
| JPS1467 | *E. coli* TOP10 pBADA-*rhlR* (I84W)/pCS26-*rhlA*-*luxCDABE*/pACYC-empty | Amp/Kan/Tet | This study |
| JPS1468 | *E. coli* TOP10 pBADA-*rhlR* (W96A)/pCS26-*rhlA*-*luxCDABE*/pACYC-*pqsE* | Amp/Kan/Tet | This study |
| JPS1469 | *E. coli* TOP10 pBADA-*rhlR* (W96A)/pCS26-*rhlA*-*luxCDABE*/pACYC-empty | Amp/Kan/Tet | This study |
| JPS1496 | *E. coli* TOP10 pBADA-*rhlR* (D81N)/pCS26-*rhlA*-*luxCDABE*/pACYC-*pqsE* | Amp/Kan/Tet | This study |
| JPS1497 | *E. coli* TOP10 pBADA-*rhlR* (D81N)/pCS26-*rhlA*-*luxCDABE*/pACYC-empty | Amp/Kan/Tet | This study |
| JPS1498 | *E. coli* TOP10 pBADA-*rhlR* (D81K)/pCS26-*rhlA*-*luxCDABE*/pACYC-*pqsE* | Amp/Kan/Tet | This study |
| JPS1499 | *E. coli* TOP10 pBADA-*rhlR* (D81K)/pCS26-*rhlA*-*luxCDABE*/pACYC-empty | Amp/Kan/Tet | This study |
| JPS1500 | *E. coli* TOP10 pBADA-*rhlR* (W108F)/pCS26-*rhlA*-*luxCDABE*/pACYC-*pqsE* | Amp/Kan/Tet | This study |
| JPS1501 | *E. coli* TOP10 pBADA-*rhlR* (W108F)/pCS26-*rhlA*-*luxCDABE*/pACYC-empty | Amp/Kan/Tet | This study |
| JPS1502 | *E. coli* TOP10 pBADA-*rhlR* (W108Y)/pCS26-*rhlA*-*luxCDABE/*pACYC-*pqsE* | Amp/Kan/Tet | This study |
| JPS1503 | *E. coli* TOP10 pBADA-*rhlR* (W108Y)/pCS26-*rhlA*-*luxCDABE*/pACYC-empty | Amp/Kan/Tet | This study |
| JPS1527 | *E. coli* TOP10 pBADA-*rhlR* (A44M)/pCS26-*rhlA*-*luxCDABE* pACYC-*pqsE* | Amp/Kan/Tet | This study |
| JPS1528 | *E. coli* TOP10 pBADA-*rhlR* (A44M)/pCS26-*rhlA*-*luxCDABE*/pACYC-empty | Amp/Kan/Tet | This study |
| JPS1529 | *E. coli* TOP10 pBADA-*rhlR* (A44W)/pCS26-*rhlA*-*luxCDABE*/pACYC-*pqsE* | Amp/Kan/Tet | This study |
| JPS1530 | *E. coli* TOP10 pBADA-*rhlR* (A44W)/pCS26-*rhlA*-*luxCDABE*/pACYC-empty | Amp/Kan/Tet | This study |
| JPS1531 | *E. coli* TOP10 pBADA-*rhlR* (G46M)/pCS26-*rhlA*-*luxCDABE*/pACYC-*pqsE* | Amp/Kan/Tet | This study |
| JPS1532 | *E. coli* TOP10 pBADA-*rhlR* (G46M)/pCS26-*rhlA*-*luxCDABE*/pACYC-empty | Amp/Kan/Tet | This study |
| JPS1533 | *E. coli* TOP10 pBADA-*rhlR* (G46W)/pCS26-*rhlA*-*luxCDABE*/pACYC-*pqsE* | Amp/Kan/Tet | This study |
| JPS1534 | *E. coli* TOP10 pBADA-*rhlR* (G46W)/pCS26-*rhlA*-*luxCDABE*/pACYC-empty | Amp/Kan/Tet | This study |
| JPS1538 | *E. coli* TOP10 pBADA-*rhlR* (T58L)/pCS26-*rhlA*-*luxCDABE*/pACYC-*pqsE* | Amp/Kan/Tet | This study |
| JPS1539 | *E. coli* TOP10 pBADA-*rhlR* (T58L)/pCS26-*rhlA*-*luxCDABE*/pACYC-empty | Amp/Kan/Tet | This study |
| JPS1540 | *E. coli* TOP10 pBADA-*rhlR* (T58I)/pCS26-*rhlA*-*luxCDABE*/pACYC-*pqsE* | Amp/Kan/Tet | This study |
| JPS1541 | *E. coli* TOP10 pBADA-*rhlR* (T58I)/pCS26-*rhlA*-*luxCDABE*/pACYC-empty | Amp/Kan/Tet | This study |
| JPS1542 | *E. coli* TOP10 pBADA-*rhlR* (T58F)/pCS26-*rhlA*-*luxCDABE*/pACYC-*pqsE* | Amp/Kan/Tet | This study |
| JPS1543 | *E. coli* TOP10 pBADA-*rhlR* (T58F)/pCS26-*rhlA*-*luxCDABE*/pACYC-empty | Amp/Kan/Tet | This study |
| JPS1544 | *E. coli* TOP10 pBADA-*rhlR* (T58G)/pCS26-*rhlA*-*luxCDABE*/pACYC-*pqsE* | Amp/Kan/Tet | This study |
| JPS1545 | *E. coli* TOP10 pBADA-*rhlR* (T58G)/pCS26-*rhlA*-*luxCDABE*/pACYC-empty | Amp/Kan/Tet | This study |
| JPS1624 | *E. coli* SM10-λ*piR* pEXG2-*rhlR* T58V | Gent | This study |
| JPS1625 | *E. coli* SM10-λ*piR* pEXG2-*rhlR* T58L | Gent | This study |
| JPS1635 | *E. coli* BL21 pETDuet-*pqsE-rhlR* | Amp | This study |
| JPS1636 | *E. coli* SM10-λ*piR* pEXG2-*rhlR* A44M | Gent | This study |
| JPS1643 | PA14 *rhlR* T58V |  | This study |
| JPS1644 | PA14 Δ*rhlI rhlR* T58V |  | This study |
| JPS1652 | PA14 *rhlR* T58L |  | This study |
| JPS1693 | PA14 *rhlR* A44M |  | This study |
| JPS1695 | PA14 Δ*rhlI rhlR* A44M |  | This study |
| JPS1771 | PA14 Δ*rhlI rhlR* T58L |  | This study |
| JPS1791 | *E. coli* BL21 pETDuet-*pqsE*-*rhlR* T58V | Amp | This study |
| JPS1792 | *E. coli* BL21 pETDuet-*pqsE*-*rhlR* L69K | Amp | This study |
| JPS1793 | *E. coli* BL21 pETDuet-*pqsE*-*rhlR* I84M | Amp | This study |
| JPS1794 | *E. coli* BL21 pETDuet-*pqsE*-*rhlR* A44M | Amp | This study |
| JPS1796 | *E. coli* BL21 pETDuet-*pqsE*-*rhlR* W108Y | Amp | This study |
| JPS1831 | *E. coli* BL21 pETDuet-*rhlR* | Amp | This study |
| JPS1832 | *E. coli* BL21 pETDuet-*rhlR* T58V | Amp | This study |
| JPS1833 | *E. coli* BL21 pETDuet-*rhlR* L69K | Amp | This study |
| JPS1834 | *E. coli* BL21 pETDuet-*rhlR* I84M | Amp | This study |
| JPS1835 | *E. coli* BL21 pETDuet-*rhlR* W108Y | Amp | This study |
| JPS1837 | *E. coli* BL21 pETDuet-*rhlR* A44M | Amp | This study |
| JPS1882 | *E. coli* DH5α pJN105 | Gent | This study |
| JPS1908 | WT PA14 attB::P*phz1A*-mScarlet | Gent | Gift from Dr. Lars Dietrich |
| JPS1909 | WT PA14 attB::P*phz2A*-mScarlet | Gent | Gift from Dr. Lars Dietrich |
| JPS1926 | PA14 Δ*rhlI* *rhlR* T58V pUCP18-p*rhlA*-mScarlet | Carb | This study |
| JPS1927 | PA14 Δ*rhlI* *rhlR* A44M pUCP18-p*rhlA*-mScarlet | Carb | This study |
| JPS1928 | PA14 Δ*rhlI* *rhlR* T58L pUCP18-p*rhlA*-mScarlet | Carb | This study |
| JPS1936 | SM10-λ*piR* pSEK109 p*phz1A*-mScarlet | Gent | Gift from Dr. Lars Dietrich |
| JPS1940 | SM10-λ*piR* pSEK109 p*phz2A*-mScarlet | Gent | Gift from Dr. Lars Dietrich |
| JPS2066 | PA14 Δ*rhlI* attB::p*phz2*-mScarlet | Gent | This study |
| JPS2189 | PA14 *rhlR* T58L attB::p*phz1*-mScarlet | Gent | This study |
| JPS2190 | PA14 *rhlR* A44M attB::p*phz1*-mScarlet | Gent | This study |
| JPS2355 | WT PA14 pJN105-6x-His-*pqsE* | Gent | This study |
| JPS2361 | PA14 *rhlR* T58L pJN105-6x-His-*pqsE* | Gent | This study |
| JPS2362 | PA14 *rhlR* A44M pJN105-6x-His-*pqsE* | Gent | This study |
| JPS2406 | WT PA14 pJN105 | Gent | This study |
| JPS2414 | PA14 *rhlR* T58V attB::P*phz1*-mScarlet | Gent | This study |
| JPS2415 | PA14 *rhlR* T58V attB::P*phz2*-mScarlet | Gent | This study |
| JPS2416 | PA14 *rhlR* T58L attB::P*phz2*-mScarlet | Gent | This study |
| JPS2417 | PA14 *rhlR* A44M attB::P*phz2*-mScarlet | Gent | This study |
| JPS2475 | *E. coli* DH5α pJN105-*pqsE* | Gent | This study |
| JPS2482 | PA14 *rhlR* T58L pJN105 | Gent | This study |
| JPS2483 | PA14 *rhlR* A44M pJN105 | Gent | This study |
