## Supplementary material for "RhlR quorum-sensing receptor ligand sensitivity regulates the differential expression of phenazine genes in *Pseudomonas aeruginosa*": Table S3

**Table S3.** Primers used in this study.

| **Oligo #** | **Sequence** | **Information** | **Purpose** |
| --- | --- | --- | --- |
| oJP166 | ttattactcgagatgaggaatgacggaggctt | pBAD-A *rhlR* forward | Restriction enzyme |
| oJP167 | ttattagaattctcagatgagacccagcgccg | pBAD-A *rhlR* reverse | Restriction enzyme |
| oJP795 | caggccgttgagcatcgccggatcca | *rhlR* I84M reverse | Mutagenic |
| oJP796 | tggatccggcgatgctcaacggcctg | *rhlR* I84M forward | Mutagenic |
| oJP797 | cgcaggccgttgagccacgccggatccacgg | *rhlR* I84W reverse | Mutagenic |
| oJP798 | ccgtggatccggcgtggctcaacggcctgcg | *rhlR* I84W forward | Mutagenic |
| oJP803 | gtggcgcacgccatacatgtagtaatcgaagcccaggcg | *rhlR* A44M reverse | Mutagenic |
| oJP804 | cgcctgggcttcgattactacatgtatggcgtgcgccac | *rhlR* A44M forward | Mutagenic |
| oJP805 | tggcgcacgccataccagtagtaatcgaagcccagg | *rhlR* A44W reverse | Mutagenic |
| oJP806 | cctgggcttcgattactactggtatggcgtgcgcca | *rhlR* A44W forward | Mutagenic |
| oJP807 | cgtgtggcgcaccatataggcgtagtaatcgaagccc | *rhlR* G46M reverse | Mutagenic |
| oJP808 | gggcttcgattactacgcctatatggtgcgccacacg | *rhlR* G46M forward | Mutagenic |
| oJP809 | gtgtggcgcacccaataggcgtagtaatcgaagc | *rhlR* G46W reverse | Mutagenic |
| oJP810 | gcttcgattactacgcctattgggtgcgccacac | *rhlR* G46W forward | Mutagenic |
| oJP1006 | gaacaggctgtcgctcgcgaccaccatttccgag | *rhlR* W96A reverse | Mutagenic |
| oJP1007 | ctcggaaatggtggtcgcgagcgacagcctgttc | *rhlR* W96A forward | Mutagenic |
| oJP1436 | aatgatacggcgaccaccgagatctacac | NextFlex forward | ChIP-seq |
| oJP1437 | CAAGCAGAAGACGGCATACGAGAT | NextFlex reverse | ChIP-seq |
| oJP1697 | agttctgcatctgggttcgctccagccaggcc | *rhlR* Y72T reverse | Mutagenic |
| oJP1698 | ggcctggctggagcgaacccagatgcagaact | *rhlR* Y72T forward | Mutagenic |
| oJP1699 | agttctgcatctgggctcgctccagccaggcc | *rhlR* Y72A reverse | Mutagenic |
| oJP1700 | ggcctggctggagcgagcccagatgcagaact | *rhlR* Y72A forward | Mutagenic |
| oJP1701 | gccatggacctcgaccttcggccgggtg | *rhlR* T58V reverse | Mutagenic |
| oJP1702 | cacccggccgaaggtcgaggtccatggc | *rhlR* T58V forward | Mutagenic |
| oJP1731 | ctggtatcgctccatccaggccttgggat | *rhlR* L69M reverse | Mutagenic |
| oJP1732 | atcccaaggcctggatggagcgataccag | *rhlR* L69M forward | Mutagenic |
| oJP1735 | tgcatctggtatcgctcatcccaggccttgggatagg | *rhlR* L69D reverse | Mutagenic |
| oJP1736 | cctatcccaaggcctgggatgagcgataccagatgca | *rhlR* L69D forward | Mutagenic |
| oJP1737 | catctggtatcgctccttccaggccttgggatag | *rhlR* L69K reverse | Mutagenic |
| oJP1738 | ctatcccaaggcctggaaggagcgataccagatg | *rhlR* 69K forward | Mutagenic |
| oJP2065 | ggatcgccggattcacggccccgta | *rhlR* D81N reverse | Mutagenic |
| oJP2066 | tacggggccgtgaatccggcgatcc | *rhlR* D81N forward | Mutagenic |
| oJP2067 | tcgcgagcctcgttgaagagcatccggctc | *rhlR* W108F reverse | Mutagenic |
| oJP2068 | gagccggatgctcttcaacgaggctcgcga | *rhlR* W108F forward | Mutagenic |
| oJP2069 | ctacggggccgtgaagccggcgatcctca | *rhlR* D81K forward | Mutagenic |
| oJP2070 | tgaggatcgccggcttcacggccccgtag | *rhlR* D81K reverse | Mutagenic |
| oJP2071 | cagagccggatgctctataacgaggctcgcgatt | *rhlR* W108Y forward | Mutagenic |
| oJP2072 | aatcgcgagcctcgttatagagcatccggctctg | *rhlR* W108Y reverse | Mutagenic |
| oJP2095 | ATTCGGTACCTTAATTAATTTCCAC | pEXG2 forward | HiFi assembly |
| oJP2096 | CCTGCAGAAGCTTGCTTTAC | pEXG2 reverse | HiFi assembly |
| oJP2139 | cttcacccggccgaagctagaggtccatggcacct | *rhlR* T58L forward | Mutagenic |
| oJP2140 | aggtgccatggacctctagcttcggccgggtgaag | *rhlR* T58L reverse | Mutagenic |
| oJP2141 | ttcacccggccgaagatcgaggtccatg | *rhlR* T58I forward | Mutagenic |
| oJP2142 | catggacctcgatcttcggccgggtgaa | *rhlR* T58I reverse | Mutagenic |
| oJP2143 | cacccggccgaagttcgaggtccatggc | *rhlR* T58F forward | Mutagenic |
| oJP2144 | gccatggacctcgaacttcggccgggtg | *rhlR* T58F reverse | Mutagenic |
| oJP2145 | cacccggccgaagggcgaggtccatggc | *rhlR* T58G forward | Mutagenic |
| oJP2146 | gccatggacctcgcccttcggccgggtg | *rhlR* T58G reverse | Mutagenic |
| oJP2181 | gtaaagcaagcttctgcaggatgaggaatgacggaggctttttg | pEXG2 *rhlR* forward | HiFi assembly |
| oJP2182 | aattaattaaggtaccgaattcagatgaggcccagcgc | pEXG2 *rhlR* reverse | HiFi assembly |
| oJP2207 | AAGCTTGCGGCCGCATAATG | pETDuet_mcs1 forward | HiFi assembly |
| oJP2208 | GTGGTGATGATGGTGATGGC | pETDuet_mcs1 reverse | HiFi assembly |
| oJP2209 | gccatcaccatcatcaccacATGTTGAGGCTTTCGGCTCC | *pqsE*_mcs1 forward | HiFi assembly |
| oJP2210 | cattatgcggccgcaagcttTCAGTCCAGAGGCAGCGC | *pqsE*_mcs1 reverse | HiFi assembly |
| oJP2211 | CTCGAGTCTGGTAAAGAAAC | pETDuet_mcs2 forward | HiFi assembly |
| oJP2212 | ATGTATATCTCCTTCTTATACTTAAC | pETDuet_mcs2 reverse | HiFi assembly |
| oJP2213 | tataagaaggagatatacatATGAGGAATGACGGAGGCTTTTTG | *rhlR*_mcs2 forward | HiFi assembly |
| oJP2214 | gtttctttaccagactcgagTCAGATGAGGCCCAGCGC | *rhlR*_mcs2 reverse | HiFi assembly |
| oJP2345 | CCGTCCTGTGAAATCTGGCA | *rhlA* forward | direct ChIP-qPCR |
| oJP2346 | TGTGTGGGTCTTGCAGATCG | *rhlA* reverse | direct ChIP-qPCR |
| oJP2349 | CACACGACTGGGTTAGGACC | intergenic forward | direct ChIP-qPCR |
| oJP2350 | ACAGGCTCGAACCAAAAGGT | intergenic reverse | direct ChIP-qPCR |
| oJP2717 | TCTTCGTGATCTGAAGCCATT | attB in check fwd | Integration check |
| oJP2718 | ctttatagatcagggtgccatcc | mS in check rev | Integration check |
| oJP2720 | AGACAGCGTAACAATCGAACG | *lecB* forward | direct ChIP-qPCR |
| oJP2721 | GACGCATCGTTCAGCCAATC | *lecB* reverse | direct ChIP-qPCR |
| oJP3133 | atatatatgctagcCCATGGGCAGCAGCCATC | *pqsE* forward pJN105 | HiFi assembly |
| oJP3134 | atatatatactagtAAGCTTTCAGTCCAGAGGCAG | *pqsE* reverse pJN105 | HiFi assembly |
